# Integrated combinatorial functional genomics and spatial transcriptomics of tumors decodes genotype to phenotype relationships

**DOI:** 10.1101/2024.05.14.593940

**Authors:** Marco Breinig, Artem Lomakin, Elyas Heidari, Michael Ritter, Gleb Rukhovich, Lio Böse, Luise Butthof, Lena Wendler-Link, Hendrik Wiethoff, Tanja Poth, Felix Sahm, Peter Schirmacher, Oliver Stegle, Moritz Gerstung, Darjus F. Tschaharganeh

## Abstract

Linking the complex genetic changes underlying cancer to relevant disease-phenotypes poses a challenge. Therefore, we present CHOCOLAT-G2P, a scalable approach that integrates multiplex in vivo functional genomics with spatial transcriptomics. By redeploying RNA-templated ligation probes of commercial spatial transcriptomics technology, we streamline mapping composite genetic alterations and transcriptome-wide phenotyping on the same tissue section on a single readout platform. Using this framework, we studied combinatorial effects of 8 perturbations that induce autochthonous mosaic liver tumors sampled from 256 genotypes. Interrogating 324 tumors across six ∼6×6 mm^2^ sections, we charted phenotypic landscapes of genotypically-defined tumor ecosystems, revealing zonation-associated hepatocellular carcinoma subclasses and associations between tumor subtypes and stromal-as well as immune-cell signatures. Further, we decoded epistasis within compound genotypes uncovering opposing roles of *Vegfa* and mutant *Ctnnb1* to cholangiocarcinoma development. Thus, CHOCOLAT-G2P lays a foundation to decipher how combinations of alterations interact to reprogram tumor cells and their microenvironment within the holistic context of tissue and whole organisms. (https://chocolat-g2p.dkfz.de/).

## MAIN

Cancer, like many other complex diseases, is caused by a combination of multiple genetic alterations^1,2^. Transitioning from portraying these genetic changes to understanding their phenotypic consequences by comparing human samples is, however, constrained by environmental influences, genetic diversity among patients, pervasive epistasis, and the complexity of multicellular tissue structure^3–5^. Consequently, it remains poorly understood how combinations of alterations reprogram cells and their interaction with the tissue environment.

Genetic screens conducted in model systems have proven valuable for decoding genotype-phenotype relationships in controlled settings^6,7^ However, current approaches for cancer-relevant in vivo functional genomics mostly investigate the effect of singular or combinatorial alterations on proliferation and tumorigenesis without considering spatial niches in which cancer cells competitively develop and grow^8–12^. Even though comprehensive studies that focus on the impact of individual alterations of tumors on their respective immunological microenvironments^13,14^ are emerging, gaps still remain in our understanding of how genetic alterations jointly rewire tumor cells and surrounding cellular ecosystems.

Here, to address this limitation, we developed CHOCOLAT-G2P (charting higher order combinations leveraging analysis of tissue to investigate genotype-to-phenotype relations, Fig. 1a), an experimental framework designed to facilitate the functional exploration of complex genotype-phenotype associations at the tissue-level (https://chocolat-g2p.dkfz.de/).

**Fig. 1:**
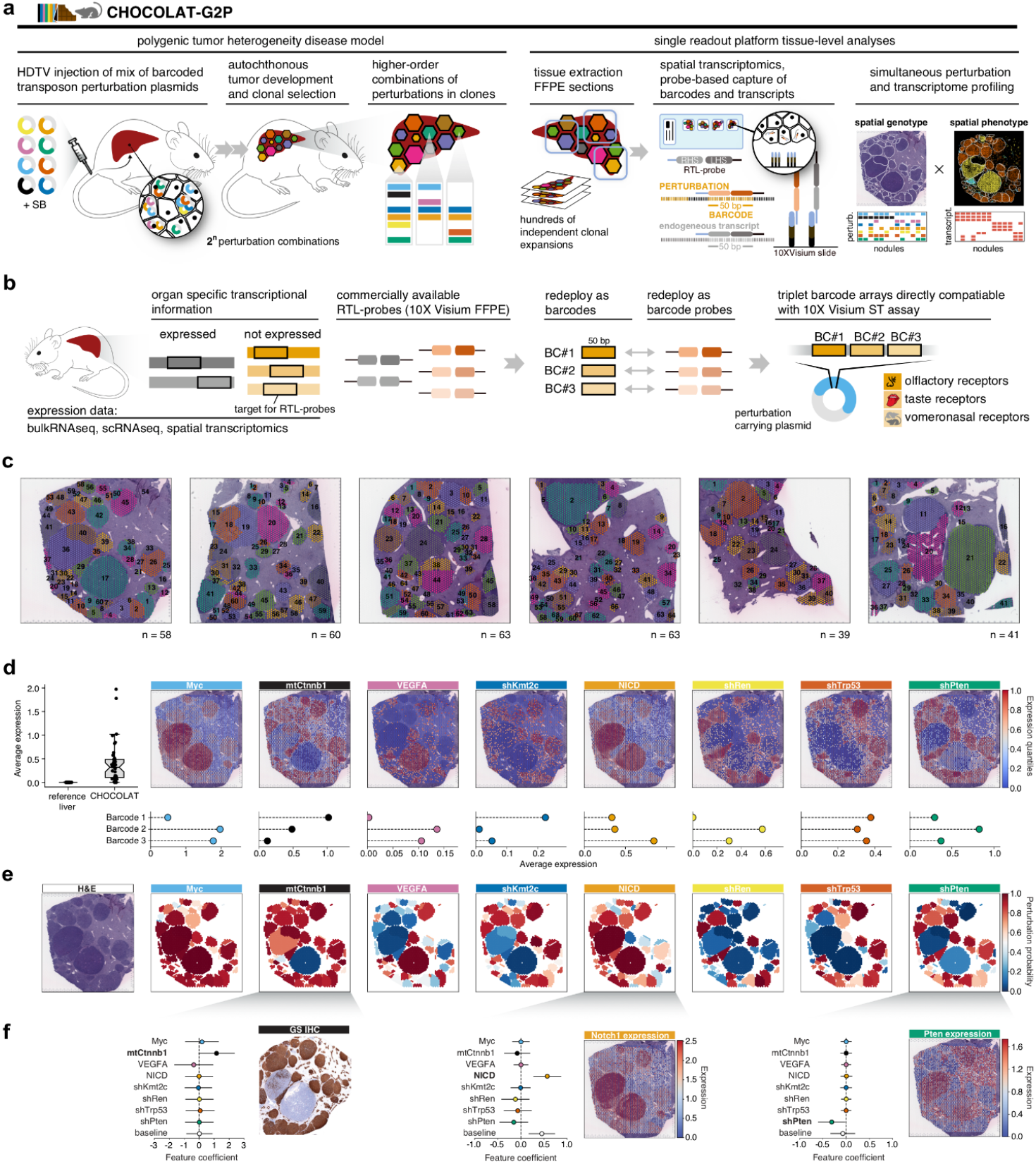
CHOCOLAT-G2P spatially resolves multiplexed genetic perturbations in hundreds of co-existing cancer clones. **a**, CHOCOLAT-G2P framework. From left: Hydrodynamic tail vein (HDTV) injection of a pool of barcoded perturbation plasmids and sleeping-beauty (SB)-transposon-mediated stable integration into the genome of hepatocytes. Higher-order combinatorial perturbations drive mosaic liver tumor development in a conceptual 2^n^ combination space for clonal selection. Direct barcode identification is achieved by linking perturbations to 50 nt barcode sequences that are captured and identified by RTL-probes as embedded in the 10X Visium for FFPE platform. Endogenous transcripts are captured alongside barcodes, hence enabling simultaneous mapping of genotypes (as defined by the presence of perturbations) and phenotypes (as defined by transcriptional signatures) on the same tissue section.**b**, Barcode selection. Transcripts not expressed in murine liver are identified using public databases. Their respective 50 nt RTL-probe capture sequences are used as barcodes detected by redeployed commercially available RTL-probes provided with the 10X Visium for FFPE mouse kit (see Methods for details). Barcodes derived from chemosensory receptor transcripts are embedded in perturbation plasmids as triplet-arrays. **c**, Establishing hundreds of co-existing tumors within native tissue context. Respective H&E-stained tissue samples for 6 topographically separated regions of ∼6×6 mm that were used for 10X Visium for FFPE-ST analysis. In total, 324 nodules (color-coded and numbered) were annotated. Colors were chosen arbitrarily. **d**, Perturbation-specific barcode identification. Left: Average log1p-transformed expression of all 38 barcode-associated transcripts used. Data combines the reference control liver datasets from^21^ and the 6 main ST-samples (CHOCOLAT-G2P) depicted in **(b)**. Panel display: Spatially resolved expression of triplet barcodes for each of the 8 perturbations. Aggregated log1p-transformed and quantile-rescaled expression per 10X Visium spot. A representative sample is shown. Lower panel: Average log1p-transformed expression of individual barcodes within each triplet array for each perturbation, averaged across the 6 ST-samples depicted in **(c). e**, Converting barcode signals to perturbation maps. Spatially resolved visualization of the inferred probabilities indicating the presence or absence of each of the 8 perturbations associated with annotated tumor nodules (Methods). A representative sample is displayed. Both the quantitative barcode expression **(d)** and probabilities **(e)** for all samples can be explored through the interactive web browser: https://chocolat-g2p.dkfz.de/. **f**, Validation of inferred perturbation integration. A generalized linear model (GLM) predicts phenotype expression signals based on the estimated probabilities of perturbation presence (Methods). Phenotypes are defined as direct target transcripts associated with perturbations such as shPten-*Pten* and NICD-*Notch1*. Expression data are log1p-transformed. GS, a well-established marker for active WNT signaling in murine livers, is used to infer mtCtnnb1-GS-positive phenotype via immunohistochemistry on a corresponding serial section. Baseline depicts background phenotype marker expression. Error-bars depict 3σ CI.

## RESULTS

### CHOCOLAT-G2P charts combinatorial genetic perturbations in hundreds of coexisting cancer clones

#### Hijacking probe-based transcript capture

CHOCOLAT-G2P merges and advances technologies, including multiplexed perturbation in vivo functional genomics, molecular barcoding and spatial omics readouts^3,5,7,9,13^, to induce and spatially map combinatorial perturbations and simultaneously characterize the resulting neoplastic phenotypes on the same sample using a single readout platform (Fig. 1a).

To illustrate CHOCOLAT-G2P’s capabilities, we modified an autochthonous murine mosaic liver cancer model to allow for the creation of co-existing, genetically-diverse tumors^7,15^. This approach relies on hydrodynamic tail vein (HDTV) injection of pooled molecular-barcoded plasmids and sleeping beauty (SB) transposon-based methods to stably integrate traceable higher-order combinations of genetic alterations in hepatocytes within their tissue context (Fig. 1a; Methods). We named this strategy RUBIX (random unique barcode integration combinatorics). In this study, we modeled complex cancer genetics by concentrating on alterations associated with liver cancer including overexpression of oncogenic drivers (*Myc*, mutant *Ctnnb1, Vegfa*, NICD) and silencing of tumor suppressors (*Trp53, Pten, Kmt2c)* with short hairpin RNA (shRNA), alongside a frequently used Renilla luciferase (shRen) targeting control construct^14,16–18^ (Extended Data Fig.1 and 3). Defined by combinations of 8 perturbations, we consequently anticipated testing 2^8^=256 possible genotypes within a single RUBIX experiment (Fig. 1a).

To simultaneously identify perturbations and obtain comprehensive tissue-level phenotypic information, our approach leverages spatial transcriptomics (ST)^5^ based on targeted transcript capture via RNA-templated ligation (RTL) probes^19^, commercially available with 10X Visium for formalin-fixed paraffin-embedded (FFPE) samples. In order to detect the introduced perturbations, CHOCOLATE-G2P uses perturbation plasmids extended with 50 nt barcodes amenable to RTL-probe capture (Fig. 1a). We named this method PERTURB-CAST (perturbation barcode capture spatial transcriptomics). To ensure immediate compatibility with default commercial kits, we redeploy 10X Visium RTL-probes capturing chemosensory receptor transcripts that are not expressed in mouse liver for barcode identification. Specifically, we exploited 50 nt RTL-probe capture sequences of olfactory-, taste-, and vomeronasal-receptor transcripts as molecular barcodes (Fig. 1b; Extended Data Fig. 2; Methods). To achieve robustness, triplet barcode-arrays were included in each perturbation construct to enable detection by three individual RTL-probes that are pre-existing components of 10X Visium spatial transcriptomics kits (Fig. 1b; Extended Data Fig. 2 and 3; Methods). In this pilot study, we distributed a total of 38 redeployed barcodes amenable to RTL-probe capture, as well as additional complementary barcodes (e.g. peptides)^13,20^ for orthogonal readouts (Extended Data Fig. 3; Methods) across 8 perturbation plasmids used for HTDV injection in a pilot RUBIX experiment.

Liver samples were collected 10 weeks after HTDV injection and processed to FFPE specimens (Extended Data Fig. 4a). For spatial transcriptomics, 6 topographically-separated regions of interest were selected based on histopathological assessment of hematoxylin and eosin (H&E) stained sections (Fig. 1c; Extended Data Fig. 4a-e). 10X Visium or 10X CytAssist were conducted for a total of 12 samples covering these 6 regions, including serial sections as technical replicates (Extended Data Fig. 4f-h). A total of 324 tumor nodules were identified across the 6 segregated sections (Fig. 1c). Of note, spatial transcriptomics helped distinguish overlapping nodules that appeared to be single lesions from the histopathological perspective (Extended Data Fig. 5).

Assessing the feasibility of our novel PERTURB-CAST barcoding strategy, we observed that most barcode signals were readily detected in the CHOCOLAT-G2P liver samples. Strikingly, barcode signals were spatially confined and closely tracking the areas of microscopic tumor nodules as revealed by H&E (Fig. 1d). In contrast, we observed that none of the 38 redeployed barcode sequences were detected in publicly available murine liver 10X Visium datasets^21^ (Fig. 1d, left). Reassuringly, with few exceptions (e.g. *Olfr1033, Olfr1358*), chemosensory receptor-transcripts that provided the repertoire for barcode redeployment (n=1216) were generally not detected by 10X Visium in murine livers (Extended Data Fig. 6a,b). Even though 5/38 redeployed barcodes showed insufficient signal across all samples investigated (average expression < 0.05, counts per 10^4^, log1p-transformed) and detection strength of individual barcodes varied, barcode-triplets enabled us to spatially identify all 8 perturbations (Fig. 1d; Extended Data Fig. 6).

### Probe-based barcode capture maps perturbations across tissue

Given the observed uncertainties related to individual barcode readouts, we used a variational Bayesian model, which accounts for multiple sources of variability including e.g. local 10X Visium spot sensitivity, to assign perturbations to each nodule (Fig. 1e; Extended Data Fig. 8; Methods). Of note, nodule-level predictions correlated across serial tissue sections and between 10X Visium and 10X CytAssist replicate experiments (Person’s r = 0.63-0.78 depending on perturbation; Extended Data Fig. 4 and 9), demonstrating quantitative reproducibility of the approach.

Investigating the expression of the genes targeted by each perturbation provides an orthogonal readout of inferred perturbation plasmid integration. Accordingly, across lesions, a generalized linear model (GLM; Methods) confirmed the expected trends of overexpression or silencing based on the corresponding perturbation, including increased expression of *Notch1* and reduced expression for *Pten* (Fig. 1f), as well as *Kmt2c* and *Trp53* (Extended Data Fig. 10b). Further validation of mtCtnnb1 was achieved by a similar analysis of glutamine synthetase (GS) immunohistochemistry, which indicates hepatic WNT/Ctnnb1-signaling activity^22^ (Fig. 1f). Lastly, the plasmids targeting *Trp53* and *Kmt2c* contained green and red fluorescent protein barcodes, respectively, providing additional immunohistochemistry-based validation (Extended Data Fig. 10c).

Thus, probe-based barcode capture spatially maps combinatorial perturbations within hundreds of co-existing tumors and provides a foundation to comprehensively chart tumor-genotypes (see interactive maps https://chocolat-g2p.dkfz.de/).

### CHOCOLAT-G2P enables tissue-level comparative genotype analyses and reveals context-dependent genetic interactions

#### Identification of cancer-drivers

Across the entirety of 324 identified nodules, the Bayesian model calculates the probabilities for all 2^8^=256 possible genotypes defined by the combinations of 8 perturbations thereby converting spatial barcode signals into genotypically-defined clonal maps (Fig. 2a; individual perturbation probabilities for 2 nodules are highlighted as an example; Extended Data Fig. 11).

**Fig. 2:**
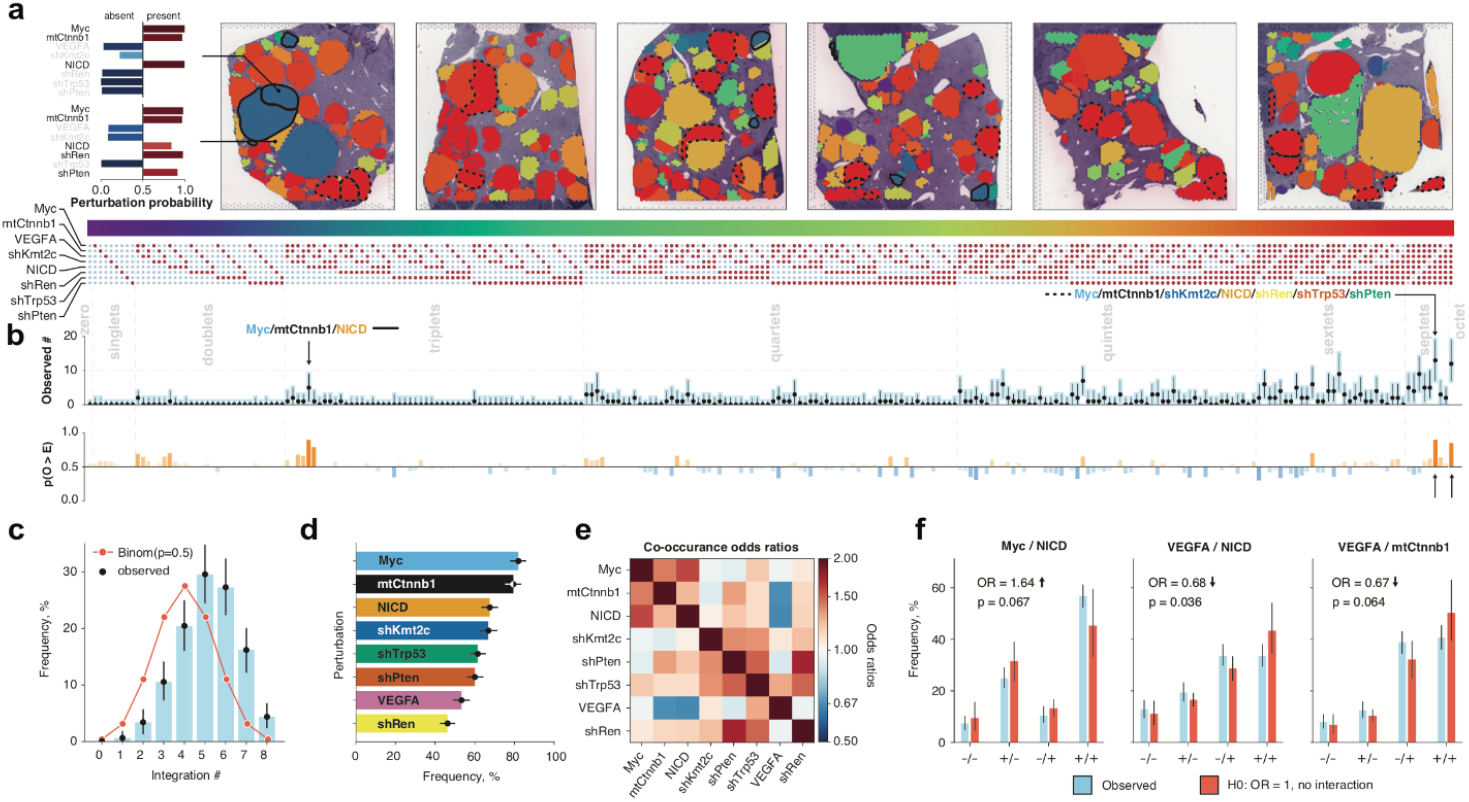
CHOCOLAT-G2P enables comparative genotype analyses and disentangles context dependent genetic interactions. **a**, Genotype maps. 2^8^ powerset embedding of spatially-mapped perturbations encompassing 324 nodules across 6 topographically separated regions. Each of the 256 combinations is color-coded. Perturbation probabilities for two representative nodules are depicted on the left. Nodules sharing representative similar genotypes are encircled and indicated in **(b). b**, Clonal selection. Upper panel: Raw, observed occurrences of genotypically-defined tumor clones (median and 95% CI) across 2^8^ powerset embedding. Highlighted genotypes are encircled in **(a)**. Lower panel: Probability p(O > E) that observed occurrences (O) deviate from the expected baseline distribution (E)(Methods). Deviations above 0.5 indicate increased tumorigenic potential (orange), values below 0.5 suggest potentially disadvantageous combinations (blue). **c**, Combinatorial order distribution. Observed distribution of the perturbation integration order (mean and 95% CI). A binomial distribution with p=0.5 is included as a reference of a random, unbiased integration rate (red line). **d**, Ranking of cancer-driving perturbations. Marginal frequencies of individual perturbations in descending order (mean and 95% CI). **e**, Pairwise co-occurrence and mutual exclusivity patterns. Odds ratios (Methods) greater than 1 suggest cooccurrence, while those less than 1 indicate mutual exclusivity. Perturbations are ordered according to **(d). f**, Identification of pairwise genetic interactions. Comparison of observed versus expected frequencies (median and 95% CI) for selected gene pairs, calculated using multiplicative models of gene interaction (Methods). Odds ratios (OR), which indicate directions of the gene interaction effect (arrows), are reported along with the corresponding p-values.

We observed that tumor clones typically exhibited combinatorial alterations with quintets being the most prevalent (∼30%; Fig. 2b,c). The absence of nodules with low integration numbers and the overall tendency towards multiple perturbations corroborates expected and previously described genetic interactions of oncogenes and tumor-suppressors, defined simply by cooperation inferred from individual experiments^16,18^. Furthermore, alterations of well-recognized oncogenes, e.g. Myc (82% of all nodules) and mtCtnnb1 (80%) occurred most frequently, while VEGFA (53%) and shRen (46%) were less frequent, possibly reflecting low tumorigenic potential (Fig. 2d).

The frequency at which perturbations are observed across nodules is dependent on the rate of successful integrations and the neoplastic potential of the combinatorial perturbation. Assuming a fixed integration rate for each perturbation allows to model an expected distribution of combinatorial events and to assess whether specific observed combinations are enriched, suggesting higher tumorigenic potential (orange) or depleted, and thus indicating disadvantageous effects (blue), independent of technical influences (Fig. 2b, bottom; Methods). Notably, among genotypes with fewer combinations, the triplet comprising Myc/mtCtnnb1/NICD emerged as frequent (n = 5, p(O > E) = 0.87) (Fig. 2b top, arrow; Fig. 2a,solid line), suggesting a strong association of this specific compound genotype with tumorigenesis. Interestingly, while septets seemed to be generally prevalent, the specific septet devoid of VEGFA (Fig. 2b top, arrow; Fig. 2a dashed line) demonstrated enrichment (n = 13, p(O > E) = 0.90) comparable to the complete octet (n = 12, p(O > E) = 0.81), whereas septets lacking e.g. mtCtnnb1 or Myc were less enriched. This in turn suggested a diminished cancer-driving effect of VEGFA in the setting of the combinatorial alterations tested.

#### Identification of co-dependencies and potential context-dependency for VEGFA

To pinpoint which perturbations contributed to the observed patterns of enriched and depleted genotypes we conducted co-occurrence odds ratio analysis. This analysis measures second-order epistatic interactions, which quantifies the deviations from purely additive effects in a commonly used multiplicative model^6,23^ (Fig. 2e,f; Supplementary Table 1; Methods). Our results revealed codependency patterns for Myc/mtCtnnb1/NICD, as well as shTrp53/shPten, aligning with previous observations^15,16,18^. The combination of Myc and NICD exhibited the most pronounced effect (OR = 1.64; pval = 0.067). In contrast, we observed a tendency towards mutual exclusivity between VEGFA and NICD (OR=0.68; pval=0.036) and between VEGFA and mtCtnnb1 (OR = 0.67; p = 0.064) (Fig. 2e,f).

Taken together these observations indicate a context-dependent oncogenic effect of VEGFA. Given VEGF’s function as a secreted molecule involved in auto/paracrine signaling^24^, we were intrigued to learn if tissue-level phenotyping of tumor ecosystems would allow us to uncover the biological underpinnings of this observation.

### CHOCOLAT-G2P maps tumor ecosystems comprising hundreds of co-existing cancer clones

#### Illuminating phenotypic landscapes

To elucidate spatial phenotypes and enable subsequent genotype-to-phenotype analyses (Fig. 3), we leveraged tissue-wide transcriptional signatures and defined sets of transcripts that characterized prevalent cell states (Extended Data Fig. 12 and 13; Methods). To finally map the complexity of tumor ecosystems, we visualized phenotype-associated transcriptional signatures within their spatial context (Fig. 3a; Extended Data Fig. 14). Further, to highlight associations for nodule-intrinsic phenotypes as well as those related to the tumor microenvironment (TME), we used transcript-resolved heatmap presentations (Fig. 3b-d). Thereby, CHOCOLAT-G2P allowed us to chart the heterogeneous phenotypic landscape of hundreds of co-existing genotypically-defined tumors and their surrounding tissue environment (see interactive maps: https://chocolat-g2p.dkfz.de/).

**Fig. 3:**
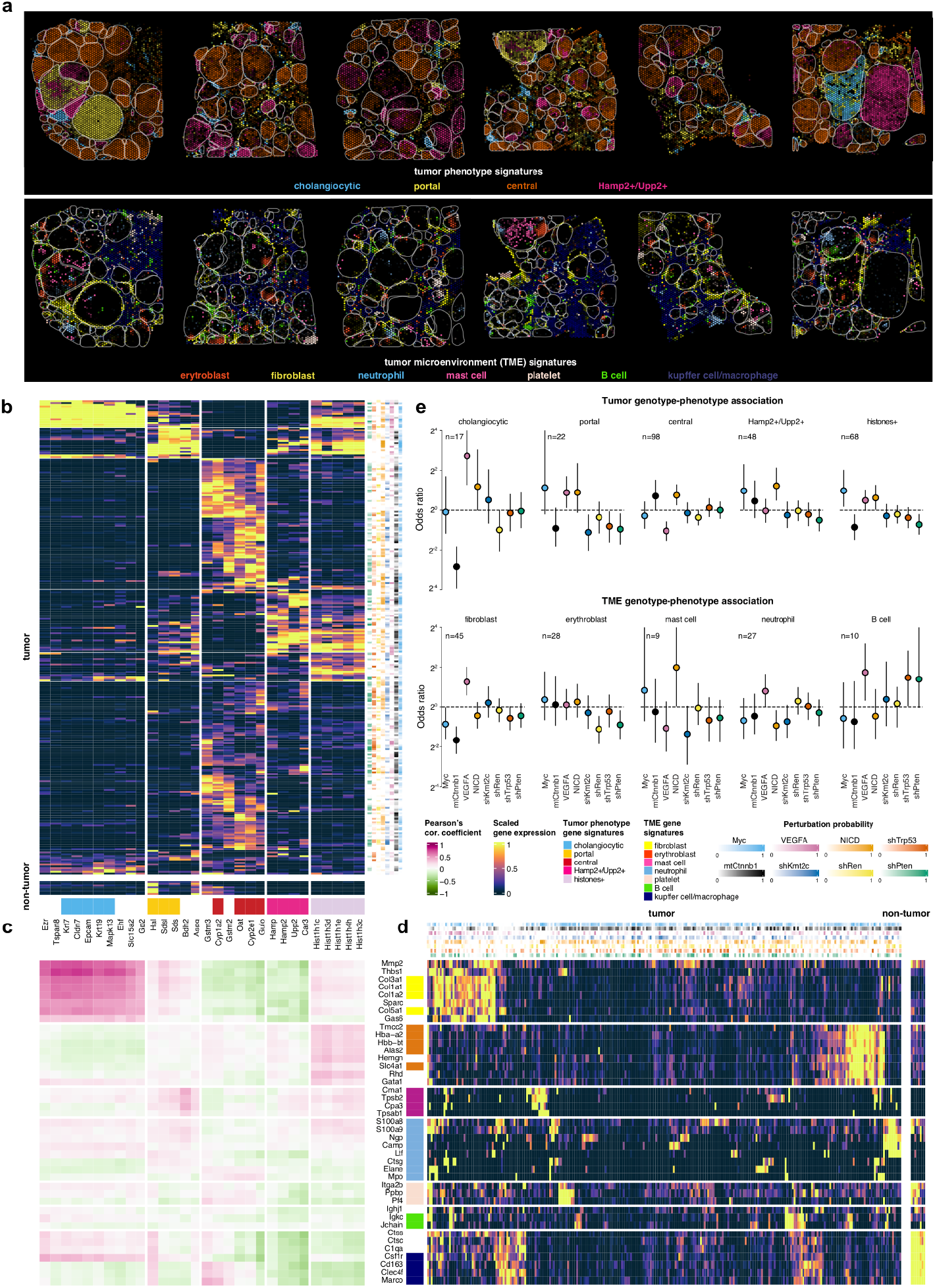
CHOCOLAT-G2P decodes relations between complex genotypes and tumor-intrinsic and microenvironmental phenotypes. **a**, Tumor ecosystem. Spatial maps of tumor-intrinsic phenotypes (top) and tumor microenvironment phenotypes (TME; bottom) across 6 topographically separated regions. Color shade depicts aggregated log1p-transformed expression of phenotype-associated transcripts (color-code as in (b) and (d)). Nodule borders are highlighted (gray). The aggregated values for all samples and underlying quantitative data of individual transcript expression can be explored through the interactive web browser interface (https://chocolat-g2p.dkfz.de/). **b**, Co-clustering of tumor-intrinsic phenotypes by associated transcripts. Tumor phenotypes are color-coded. Clustering based on Spearman correlations. Phenotypes are subdivided using hierarchical clustering. Right-hand side heatmap illustrates scaled (p^10) estimated plasmid probabilities per nodule. **c**, Associations between tumor-intrinsic and TME phenotypes. Pearson’s correlation coefficient for each pair of tumor-intrinsic and TME phenotype associated transcripts across all nodules. **d**, Co-clustering of TME phenotypes by associated transcripts. TME phenotypes are color-coded. Clustering based on Spearman correlations. Phenotypes are subdivided using hierarchical clustering. Top heatmap illustrates scaled (p^10) estimated plasmid probabilities per nodule as in **(b). e**, Identification of genotype-phenotype relations. Odds ratios comparing the prevalence of perturbations between phenotypic groups and the remainder of the nodules (total n=324) for tumor intrinsic phenotypes (top) and tumor microenvironment (bottom). OR > 1 indicates enrichment of perturbations within the phenotypic group; OR < 1 indicates depletion. Number of nodules with a given phenotype (n) is indicated. Note that groups are not mutually exclusive. Median and 90% CIs are reported.

#### Stratification of liver tumor subtypes

Our approach readily distinguished prominent subtypes of liver tumors (Fig. 3a upper panel). First, we observed nodules with cholangiocyte-like transcriptional signatures (e.g.*Krt19+/Cldn7+*) indicative of cholangiocarcinoma (CCA)^25,26^ (Fig. 3a,b; Extended Data Fig. 14a). Indeed, microscopic inspection classified these tumor nodules as cholangiocarcinomas exhibiting a glandular growth pattern and stroma deposition as well as CK19-protein expression by immunohistochemistry (Extended Data Fig. 15). Cholangiocarcinoma is the second most common type of liver cancer next to hepatocellular carcinoma (HCC) and both tumor types can develop from hepatocytes in the HDTV injection-based mouse model^25,26^. Interestingly, CHOCOLAT-G2P pinpointed, among others, expression of *Slc15a2* (Solute Carrier Family 15 Member 2) for which genomic variants have been linked to sorafenib-therapy response^27^, as well as *Gp2* (Pancreatic Glycoprotein 2) as being associated with cholangiocarcinoma (Fig. 3b; Extended Data Fig. 14a). The latter observation harmonizes with earlier research, suggesting that anti-GP2 IgA autoantibodies enable early cholangiocarcinoma detection in subsets of human patients^28^. Moreover, we could immediately relate the prominent second and third cluster of tumor nodules to metabolic liver zonation. Spatial division of metabolic functions is not only central to liver tissue organization under physiological conditions^29,30^ but has been proposed to enable molecular classification of human hepatocellular carcinoma^31^. In alignment with this zonation-based molecular classification, CHOCOLAT-G2P enabled us to stratify nodules either as portal-like (e.g. *Sds+/Sdsl+*) or central-like (e.g. *Cyp2e1+/Oat+*)^29,30^ (Fig.3 a,b; Extended Data Fig.14b,c). Lastly, focusing on the tumors that could not readily be assigned to the aforementioned subtypes, we identified a fraction of nodules that revealed enrichment of *Hamp, Hamp2* (hepcidin antimicrobial peptide) and *Upp2* (Uridine Phosphorylase 2) expression (Fig. 3a (*Hamp2+/Upp2+*); Extended Data Fig. 14d). *Hamp* and its paralog *Hamp2* have both been associated with midlobular zonation^30^, a feature of liver structure important for regeneration^32^. *Upp2*, on the other hand, is involved in pyrimidine salvage, which fuels glycolysis and enables growth of cancer cells under nutrient-limited conditions^33,34^. We further identified subgroups of nodules sharing cholangiocytic as well as portal-like features (Fig. 3b), an observation in agreement with a proposed hybrid periportal hepatocyte cell type^35^. Likewise, subsets of nodules from the major classes, with the exception of central-like nodules, shared striking enrichment of numerous histone-associated transcripts (Fig. 3b). Upregulation of genes encoding histone proteins is described as the most prominent gene regulatory program at the G1/S-phase transition in human pluripotent cells^36^.

#### Tumor-stroma and tumor-immune cell connections

Next, by focusing on cellular ecosystems of the liver tumor microenvironment (Fig. 3a lower panel (TME); Fig. 3d), we identified fibroblast-associated transcriptional signatures (e.g. *Col1a1+/Col3a1+*)^37,38^ prominent at the tumor-stroma border. We further observed regionally-segregated expression patterns associated with hematopoietic/immune cell clusters (Fig. 3a). These included signatures likely associated with erythroblasts (e.g. *Hbb-bt+/Slc4a1+*)^39,40^, platelets (e.g. *Pf4+/Itga2+*)^41^, mast cells (e.g. *Cpa3+/Cma+*)^42,43^, B-cells (*Jchain+/Igkc+)*^44^, and neutrophils (e.g. *Elane+/Mpo+*; *Ngp+/Camp+*)^40,45,46^. Signatures associated with Kupffer cells/macrophages (e.g. *Marco+/Clec4f+; Csf1r+/C1qa+*)^3,21^ were primarily detected within the non-tumor compartment (Extended Data Fig. 14f-k).

Our approach immediately revealed connections between tumor-intrinsic cell states and the microenvironment, such as a notable link between cholangiocarcinoma and fibroblast-like signatures (Fig. 3c). This observation aligns with data indicating that cancer-associated fibroblasts (CAFs) are the major cellular component of human cholangiocarcinoma-associated desmoplastic stroma^38^. Indeed, unbiased phenotyping grouped fibroblast-like signatures alongside *Gas6* (Growth Arrest-Specific 6) and *Thbs1* (Thrombospondin 1), both of which were identified as marker transcripts for a mechanoresponsive CAF subpopulation^37^ (Fig. 3b; Extended Data Fig. 14f). Our results further pointed towards additional associations, such as a link between cholangiocarcinoma and macrophages (e.g. *Csf1r+/C1qa+*), as well as a connection between enriched erythroblast (*Hbb-bt+/Slc4a1+*) occurrence and the histone-associated subgroup of nodules (Fig. 3c).

### CHOCOLAT-G2P decodes relations between complex genotypes and tumor-intrinsic and micro-environmental phenotypes

#### Inferring genotype-relations from defined phenotypes

Using spatial maps that combine phenotypical and genotypical data from the same tissue sections enables detailed investigation of phenotype-genotype relations. (Fig. 2a; Fig. 3a-d). We therefore assigned binary phenotype labels to nodules (Extended Data Fig.16; Methods) and calculated odds ratios to assess each perturbation’s connection to specific tumor-intrinsic and micro-environmental phenotypic groups (Fig. 3e). Remarkably, within the setting of combinatorial perturbations tested, our findings indicated that cholangiocarcinoma reveal a strong positive association with VEGFA, exceeding any other observed linkage, as well as a strong negative association with mtCtnnb1 (Fig. 3e). Further, consistent with the central role of WNT-signaling in liver zonation^47^, portal-like tumors revealed negative associations whereas central-like nodules revealed positive associations with mtCtnnb1 (Fig. 3e), albeit the differential contribution of additional perturbations such as Myc demand further investigation. Focussing on relationships between tumor-genotypes and TME phenotypes, we observed strong positive associations for VEGFA with fibroblast-signatures, alongside a negative association with mtCtnnb1 (Fig. 3e), largely resembling the patterns observed for cholangiocarcinoma and being in line with their prominent spatial association (Fig. 3a,b). A similarly positive association for VEGFA was also observed for B-cell-like signatures, alongside negative association for Myc and mtCtnnb1 and positive associations for shTrp53 and shPten (Fig. 3e). Reflective of our findings, immune cell infiltration as evaluated by CD45-positive immunohistochemistry was reported in a compound Trp53/Pten knockout HDTV injection model of liver cancer^18^, whereas immune cell exclusion has been observed in a corresponding Myc/mtCtnnb1 model^16^.

#### Inferring phenotype-relations from defined genotypes

To broaden our analysis beyond binary phenotypes, we leveraged spot-level continuous expression of phenotype-associated marker transcripts. We therefore calculated associations for each perturbation using the aforementioned GLM-analyses (Fig. 17a,b). Despite its limited sensitivity to identify associations for transcripts that reveal sparse spatial expression (Methods) this analysis supported the identified associations for VEGFA and mtCtnnb1 for cholangiocyte-associated transcripts (e.g. *Krt19* and *Cldn7*). Of note, GLM-analyses uncovered additional transcript-specific contributions of perturbations not readily apparent using the nodule phenotype-binarization approach (Fig. 3e). For example, we observed a strong relation of cholangiocyte-associated transcripts *Krt19* and *Gp2* with shKmt2c perturbation (Extended Data Fig. 17a). Since multiple transcripts associated with predefined phenotype signatures shared similar ‘GLM-patterns’ (Extended Data Fig. 17a,b), we finally interrogated perturbation-phenotype associations on a transcriptome-wide scale (Methods). Focusing on 1,283 genes that showed significant associations with perturbations, we observed clusters of transcripts that correspond to specific cell states (Extended Data Fig. 17c and Supplementary Table S2). For example, global-GLM analysis grouped together fibroblast-associated transcripts such as *Col1a1, Col1a2, Gas6* and *Thbs1* or transcripts related to the aforementioned histone-enriched phenotype. Likewise, GLM-analysis aggregated *Krt19, Epcam, Gp2*, and *Krt7* all of which defined the cholangiocytic phenotype that was linked to positive associations with VEGFA and negative associations with mtCtnnb1 (Extended Data Fig. 17c, Fig.3 b,e).

## DISCUSSION

Here, we introduced CHOCOLAT-G2P, an experimental framework that seamlessly integrates higher-order combinatorial perturbation in vivo functional genomics with spatial transcriptomics interrogation by redeploying RNA-templated ligation probes from commercial technology for molecular barcode identification. Combined with the ability to induce hundreds of tumors, each with distinct combinations of alterations, in a single tissue, CHOCOLAT-G2P’s capability to characterize tumor gene expression and cellular microenvironments helps address the long-standing question how multiple genetic changes interact to shape disease-phenotypes within cellular ecosystems.

By applying CHOCOLAT-G2P to an autochthonous mouse model of liver cancer, we explored a wide range of cancer-driving combinatorial genotypes sampled from nearly all of the 2^8^ combinations of perturbations present within hundreds of co-existing tumors. The integration of spatial transcriptomics into CHOCOLAT-G2P enabled simultaneous mapping of each nodule’s genotype alongside tumor-intrinsic and microenvironment-related phenotypes on the same tissue sample. Our novel PERTURB-CAST approach eliminated the need for laborious and potentially biased complementary readouts, as well as the requirement for analyses on serial tissue samples, thereby preserving spatial relationships and providing the basis for detailed genotype-phenotype analyses (https://chocolat-g2p.dkfz.de/).

Interrogating 324 liver tumor nodules from a single experiment revealed mutual exclusivity between mutant Ctnnb1 and VEGFA, indicating epistatic fitness effects of these two alterations. Their exclusivity was further underscored by phenotypic divergence. Specifically, mutant Ctnnb1, which we identified as a crucial contributor to overall liver tumor occurrence, masked the emergence of the cholangiocarcinoma subtype, thus exemplifying Bateson’s classical definition of epistasis^23^. In contrast, VEGFA induced a cholangiocytic histology and gene expression profile (Extended Data Fig. 18). VEGFA perturbed nodules also exhibited a greater abundance of cancer associated fibroblasts compared to nodules with mutant Ctnnb1, indicating that genetic alterations also shape, and possibly coopt, their tumor microenvironment.

CHOCOLAT-G2P can be applied and extended in a number of ways. First, it is straightforward to exchange the alterations and perform combinatorial screens involving other cancer drivers^2,48^, potentially including sequential introduction of perturbations, e.g. by means of inducible expression^49^. Such RUBIX screens may also be conducted in different mouse strains or growth conditions to model the interactions between tumor genomes and host genetics, immunocompetence, environmental exposures and therapeutic interventions^7,14,50^. Second, we envision the applicability of CHOCOLAT-G2P beyond liver cancer. Currently available autochthonous animal disease models that similarly build on stable integration of perturbations include diverse tumor types such as lung, pancreas, stomach or soft tissue cancers^7,51,52^. Further, CHOCOLAT-G2P’s RTL-probe-based readout may be used with other spatial transcriptomics platforms that provide higher resolution and should enable immediate implementation of PERTURB-CAST into 10X Visium HD^53–55^. Moreover, our currently employed perturbation plasmids (Extended Data Fig. 3) are expected to be directly compatible with orthogonal barcode mapping strategies^13,20,56,57^ and only minor adjustments to match the interrogated sequences are required to use CHOCOLATE-G2P in conjunction with imaging-based platforms^58,59^. Since multiple spatial omics technologies hinge on the availability of robust RNA-detection probes, a requirement not always satisfied as evidenced by the inefficacy of PERTURB-CAST to detect 5/38 barcodes tested in this study, prospective massively parallel assays^54^ based on CHOCOLAT-G2P could be leveraged to screen for reliable probes.

In summary, CHOCOLATE-G2P provides a new multiplexed approach for higher-order combinatorial cancer screens in a single mouse. Its design, which utilizes off-the-shelf spatial transcriptomics protocols, facilitates comprehensive readouts of tumor genotypes and spatial phenotypes from the same tissue sample.

As CHOCOLATE-G2P may be extended to other alterations^57,60,61^, disease models^3,7,12,52,62^ and spatial omics technologies^5,13,59^, we envisage a broad range of applications to finally decode the relationships between complex genotypes and phenotypes.

## Data availability

We have launched a web-browser for interactive data analyses (https://chocolat-g2p.dkfz.de/).

We deposited data related to this manuscript to https://zenodo.org/records/10986436/.

## Code availability

Scripts and custom code for data analysis related to this manuscript are available at https://github.com/gerstung-lab/CHOCOLAT-G2P/.

## Author contributions

M.B. conceptualized the project, designed the methodology and experiments.

E.H. and A.L. performed 10X Visium data preprocessing.

A.L. implemented ST-based genotyping with input from E.H., M.B, O. St. and M.G.

E.H. carried out ST-based phenotyping with input from A.L., M.B, O. St. and M.G.

A.L. and E.H performed statistical analyses with input from O. St. and M.G.

L.Bö., L.Bu., L.W-L, cloned plasmids, performed animal experiments and immunohistochemistry with help from M.B.

M.R. performed 10X Visium spatial transcriptomics and NGS with help from M.B. P.S., D.F.T., and F. S. provided reagents.

T.P. supervised technical processing of FFPE liver samples.

H.W. and D.F.T. performed histopathological analysis.

H.W. and M.B. performed spatial transcriptomics-guided nodule annotation.

M.B. performed initial bulk RNAseq data analysis for probe redeployment.

M.B., A.L. and E.H. created Figures with input from D.F.T, O. St. and M.G.

A.L. and G.R. developed the dedicated webtool interface with input from E.H. and M.B.

M.B., A.L. and E.H. wrote the manuscript with input from D.F.T and M.G.

## Acknowledgements

We thank all members of the Tschaharganeh, Sahm, Gerstung and Stegle lab.

We thank the German Cancer Research Center Central Animal Laboratory, the Center for. Model Systems and Comparative Pathology of the Institute of Pathology Heidelberg, and the Tissue Bank of the National Center for Tumor Diseases for excellent technical support.

D.F.T was supported by the European Research Council (ERC Starting Grant “CrispSCNAs” (grant number: 948172) and by the Deutsche Forschungsgemeinschaft (DFG) (TS 293/3-1).

