## Extended data for "Integrated combinatorial functional genomics and spatial transcriptomics of tumors decodes genotype to phenotype relationships"

### SUPPLEMENTARY FILES:

### EXTENDED DATA FIGURES

### ONLINE METHODS

### METHODS REFERENCES

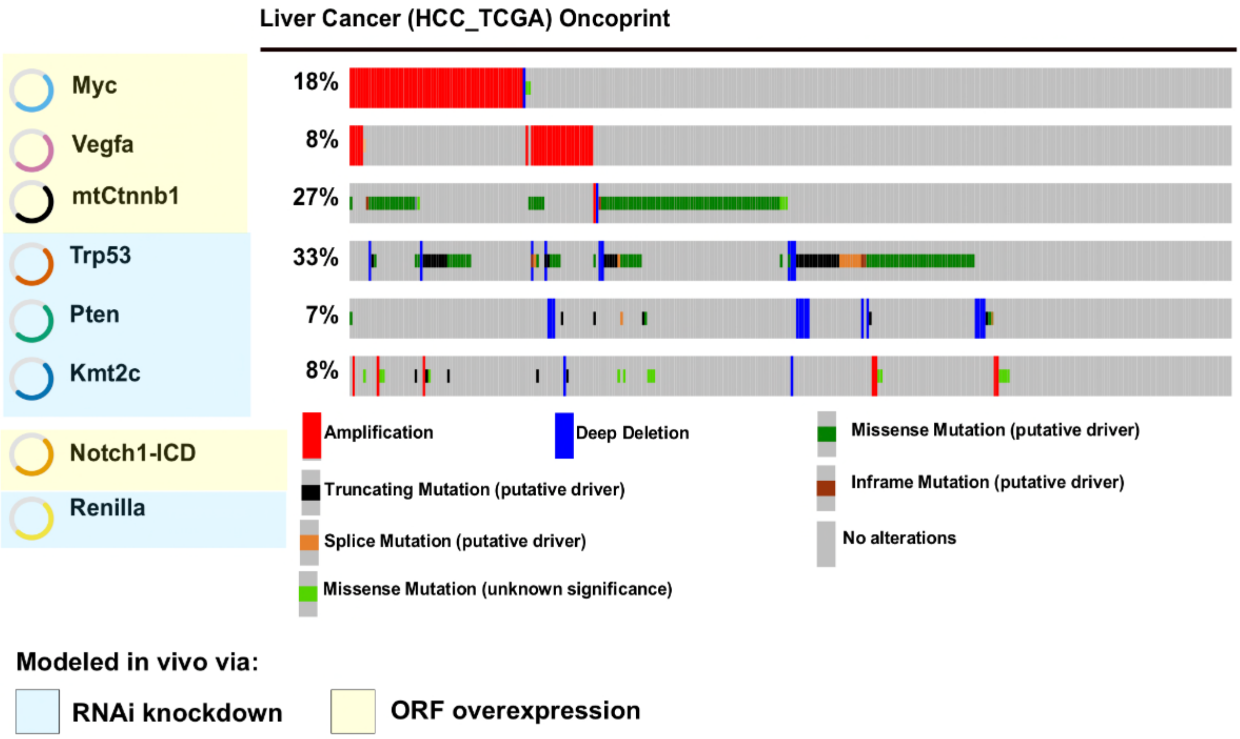


**Extended Data Fig. 1: Selection of liver-cancer relevant perturbations**

Frequent alterations observed in human liver cancer are "geno-copied" in a CHOCOLAT-G2P mouse model. (Oncoprint representations; data based on https://www.cbioportal.org/study/summary?id=lihc_tcga). Instead of solely investigating complete loss-of-function phenotypes via CRISPR/Cas knockout[13] and to avoid known complications, such as gross chromosomal rearrangements of multiplex CRISPR/Cas-knockout experiments in our mouse model[8], we aimed to employ established alternative orthogonal genetic tools[15]. Specifically, we used forced constitutive overexpression to model frequent gains/amplifications (e.g. Myc, Vegfa) as well as oncogenic mutations (e.g. mutant Ctnnb1) and RNA interference-mediated knockdown via short-hairpin (sh)RNA to model loss-of-function or loss-of-heterozygosity (e.g. Trp53, Kmt2c, Pten) in the mouse model employed. Further, we used overexpression of a notch intracellular domain (NICD), a perturbation previously used to phenocopy constitutive Notch signaling activity in the mouse model used[15]. We also used a Renilla luciferase targeting shRNA which is frequently employed as a neutral control perturbation in the mouse model used[15].

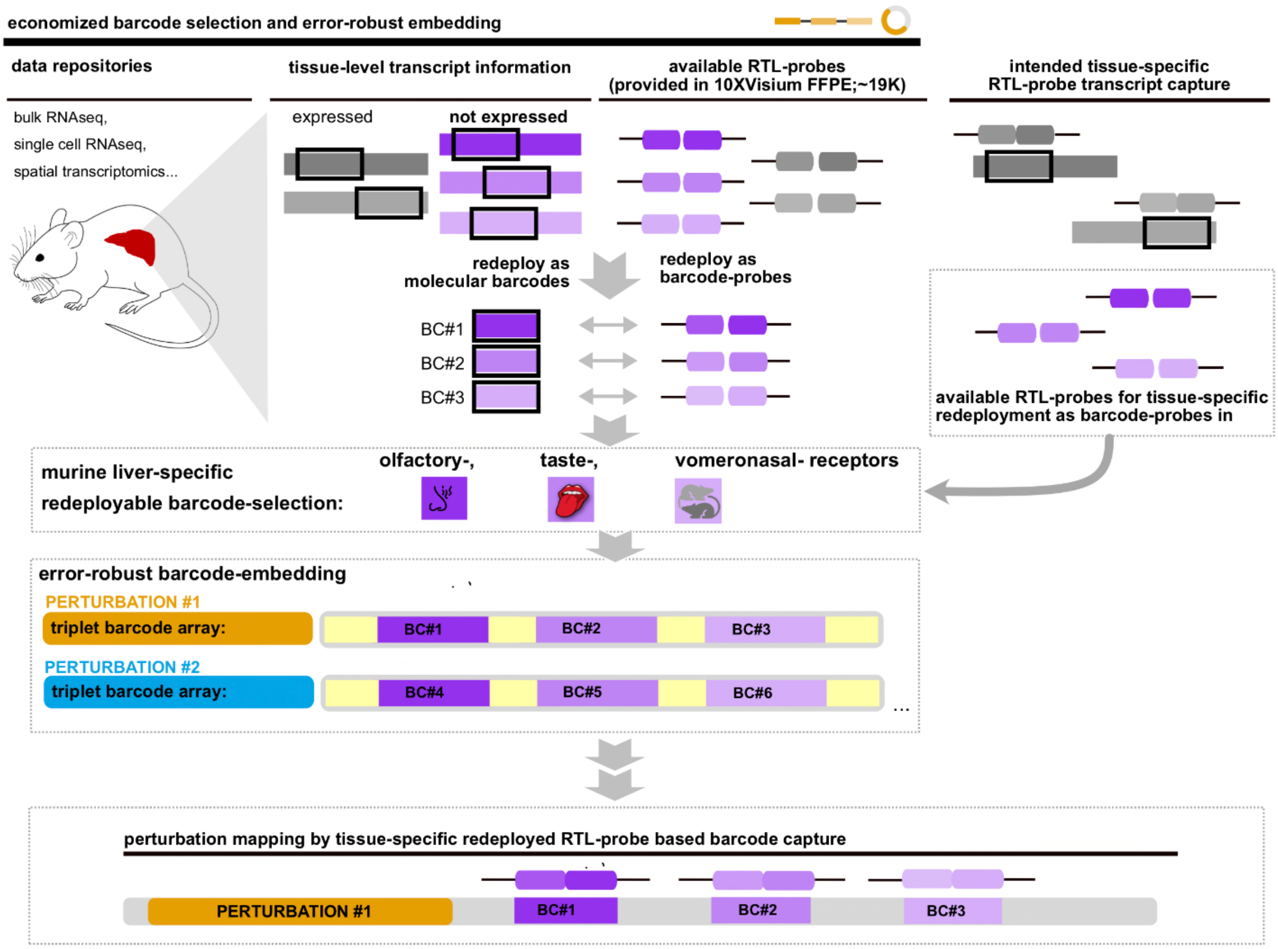


**Extended Data Fig. 2: Economized barcode selection and error-robust embedding**

Transcripts not expressed in tissue-of-interest (here murine liver) are identified using public databases. Their respective 50 nt RTL-probe capture sequences (available from 10X Genomics) are redeployed as detectable barcodes using commercially available RTL-probes against endogenous transcripts (provided with the 10X Visium for FFPE mouse kit; see Methods for details).

Murine liver-specific redeployed barcodes are derived from olfactory-, taste-, and vomeronasal-receptor transcripts.

Identified redeployable barcode sequences (Methods) are integrated into perturbation plasmids into triplet arrays. Consequently, three available RTL-probes enable identification of a single perturbation providing error-robustness. This means that if one barcode/RTL-probe pair proves non-functional, the presence of the other two sequences in the set ensure accurate identification of the perturbation. If two barcode/RTL-probe pairs prove non-functional, the presence of one sequence in the set ensures accurate identification of the perturbation. Only if three barcode/RTL-probe pairs prove non-functional, perturbation identification is unsuccessful.

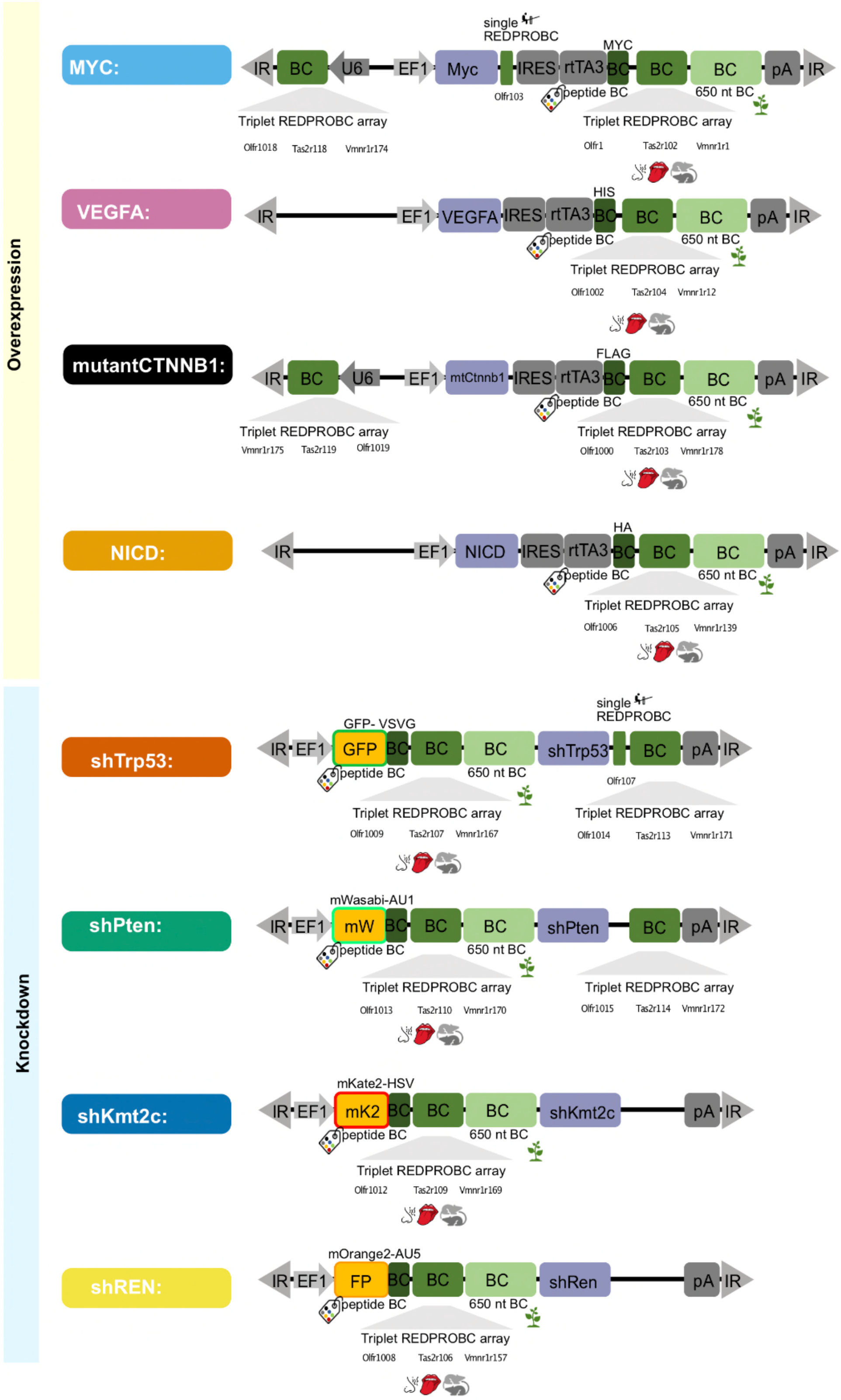


**Extended Data Fig. 3: Overview of perturbation plasmid design and molecular barcoding**

Schematic overview of Sleeping-Beauty transposon perturbation plasmids to ectopically overexpress genes-of-interest (oncogenic-driver perturbations) or shRNA to enable gene knockdown (tumor-suppressor perturbations).

Functional elements are highlighted.

IR: Inverted/direct repeats of sleeping beauty transposon; EF1: Polymerase II promoter; U6: Polymerase III promoter; pA: polyadenylation signal; IRES: internal ribosome entry site, for polycistronic expression; rtTA3: reverse tetracycline transactivator as used for the “Tet-0n system”, which is a doxycycline-inducible gene expression system. Functionality was not tested in this study.; sh: short hairpin RNA embedded in miRE context; BC: barcode.

Note that plasmids were equipped with multiple barcodes at varying positions.

Peptide BC: a barcode encoding for peptides/proteins (such as FLAG, AU1, GFP etc. as indicated). Functionality was not tested comprehensively in this study.

650 nt BC: a “long” RNA-barcode (stretches of at least 650 nts derived from the combination of multiple oligo-miner probe sequences designed against Arabidopsis thaliana Chr1). Functionality was not tested in this study.

Single REDPRO-BC: a single 50 nt barcode amenable to redeployed RTL-probe capture. Note that we used Visium Mouse Transcriptome Probe Set v1 to derive barcodes.

Triplet REDPRO-BC array: a barcode in which 3 redeployed RTL-probe capture sequences (as indicated) are embedded. Note that we used Visium Mouse Transcriptome Probe Set v1 to derive barcodes. Each 50 nt barcode is separated and flanked by ca. 20 nt spacer sequences to avoid potential steric hindrance during hybridization. Spacer sequences used were derived from T7 and T3 promoters and/or AsCas12a-DR sequences and/or 10X Capture sequences cs1 and cs2 (not shown). Functionality of spacer sequences was not tested in this study.

See Methods for further information.

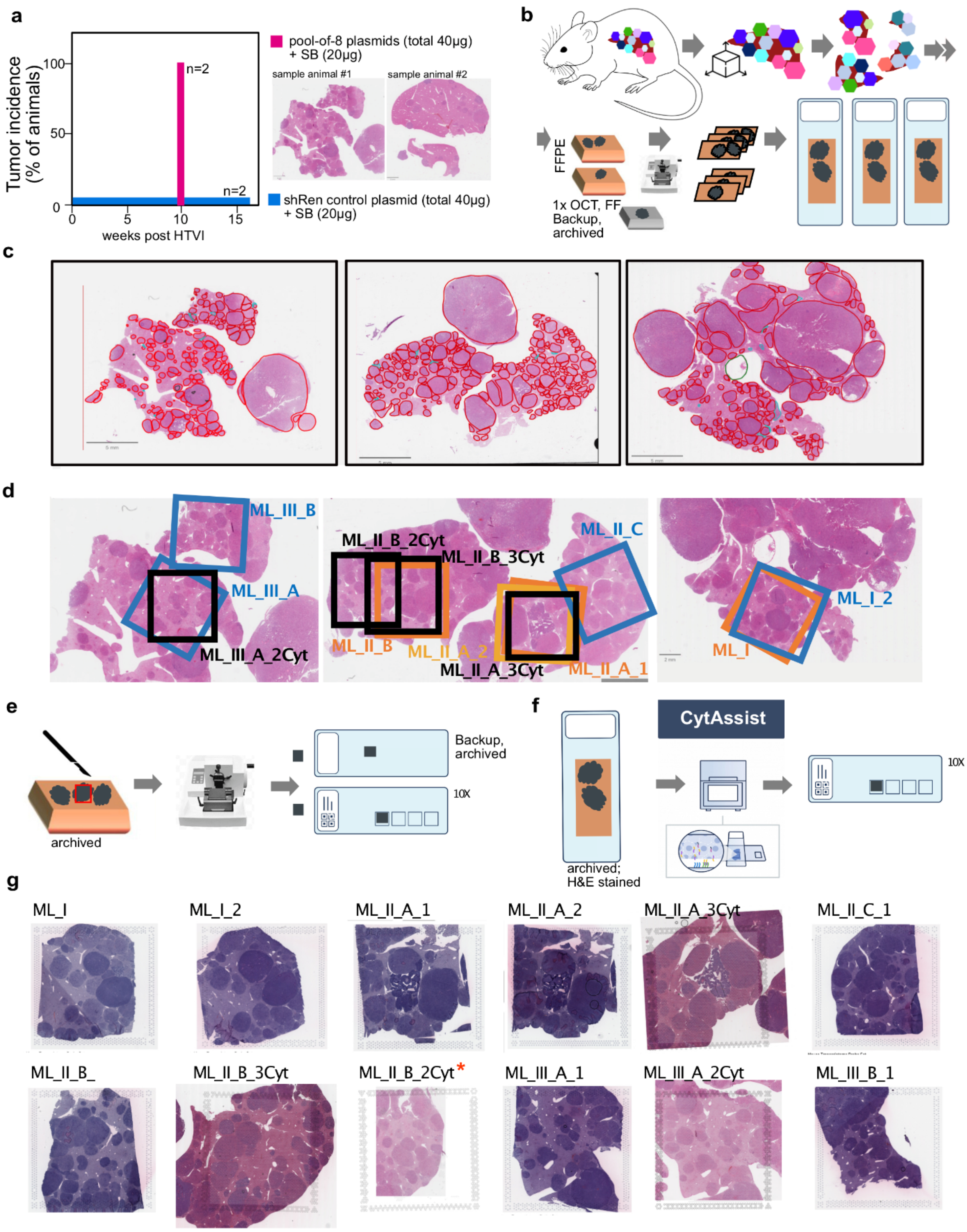


**Extended Data Fig. 4: Sample processing and ROI selection for spatial transcriptomics**

**a, Multiplexing with a pool-of-8 plasmid mix results in rapid liver tumor development.** Injection of a shRNA targeting Renilla (shRen; matching total plasmid concentration for pool-of-8 mix) served as control, n=2 each group. Representative H&E stained samples revealing multiple tumor nodules from the 2 individual animals are shown. Absence of tumors in the control group (shRen only) indicates that random integration of transposon plasmids itself is unlikely to contribute to tumorigenesis.

**b, Tissue preprocessing.** Following liver tumor development, livers were extracted, divided and processed to FFPE as well as fresh frozen specimens. FFPE samples were initially sectioned to enable sample selection.

**c, CHOCOLAT-G2P liver samples.** Overview of 3 representative FFPE samples used in this study. A total of 513 tumor nodules (red outline) were identified based on histopathological examination (based on H&E).

**d, Overview of ROIs selected for 10X Visium**. 6 segregated regions were selected across 3 FFPE samples. Squares indicate approximate position of ROIs selected for ST. Orange: first 10X Visium run, light orange: first 10X Visium run, replicate ROI; blue: second 10X Visium run; black: 10X Visium CytAssist run. Note overlap between ROIs, where serial sections are used for 10X Visium.

**e, 10X Visium workflow.** Samples for ST are derived directly from FFPE blocks and mounted on 10X Visium slides.

**f, 10X Visium CytAssist workflow**. Samples for ST are derived from sections already mounted on glass slides and transferred to 10X Visium slides using the 10X CytAssist instrument.

**g, Overview of all samples used for ST.** 12 samples were used for 10X Visium in this study. Respective H&E stainings are depicted. Note that the utility of sample ML-II_B_2Cyt is constrained by tissue detachment of the sample during the processing for 10X Visium CytAssist and was not included for further analyses.

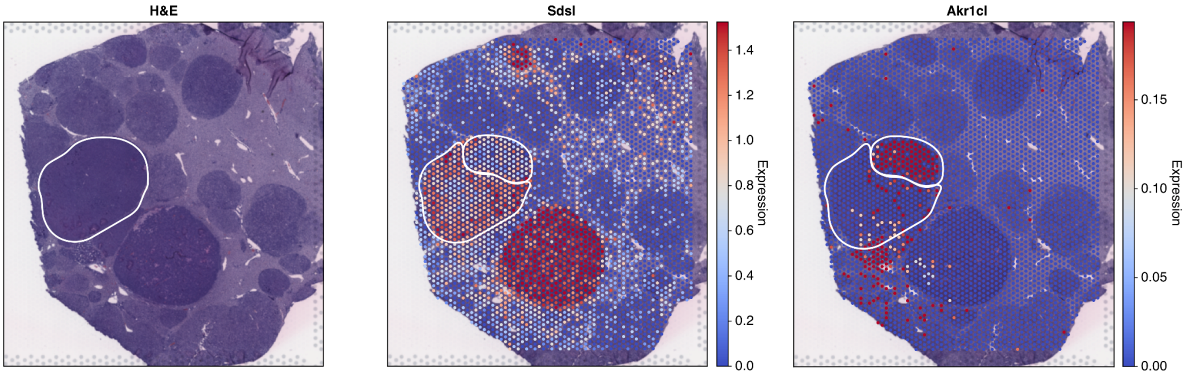


**Extended Data Fig. 5: ST enables identification of overlapping nodules**

H&E based annotation identified a single large tumor (left; encircled). ST differentiated this tumor into two nodules. To highlight example transcripts that enable differentiation, *Sdsl* is expressed in the larger tumor nodule, whereas *Akr1cl* is expressed in the smaller nodule.

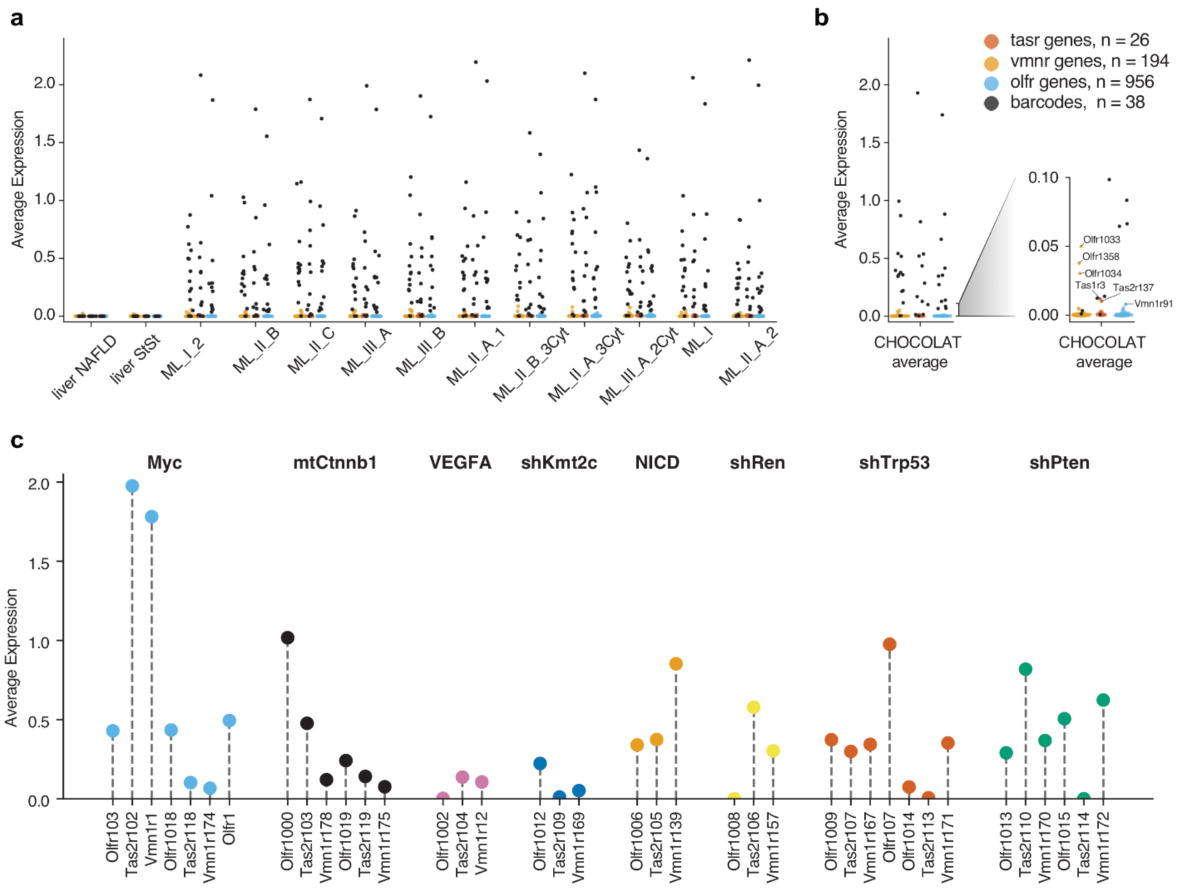


**Extended Data Fig. 6: Specific detection of redeployable barcodes by available 10X RTL-probes**

**a, Expression of all potentially redeployable barcodes derived from sensory receptor transcripts in murine liver.** Transcripts associated with three groups of sensory receptors - *Tasr* (red; n=26), *Vmnr* (yellow; n=194), *Olfr* (blue; n=956). Redeployed barcodes (black) are well separated from endogenous sensory receptor transcripts (total n=1216). Data for all 11 CHOCOLAT-G2P samples generated in this study is depicted.Reference control liver datasets (NAFLD, StSt) are from[17].

**b, Sensory receptor-associated transcripts expressed in murine liver.** Sensory receptor-associated transcripts that reveal average expression between < 0.01 and > 0.008 in murine liver are depicted. Expression values are aggregated across all 11 CHOCOLAT-G2P samples shown in (**a**).

**c, Expression of all 38 redeployed barcodes used in this study**. Grouped according to associated perturbation (see Extended Data Fig.3, Methods). Expression is averaged across 6 primary CHOCOLAT-G2P samples. Note that 5/38 revealed insufficient signal (average expression < 0.05); In all panels, expression values are log1p-transformed.

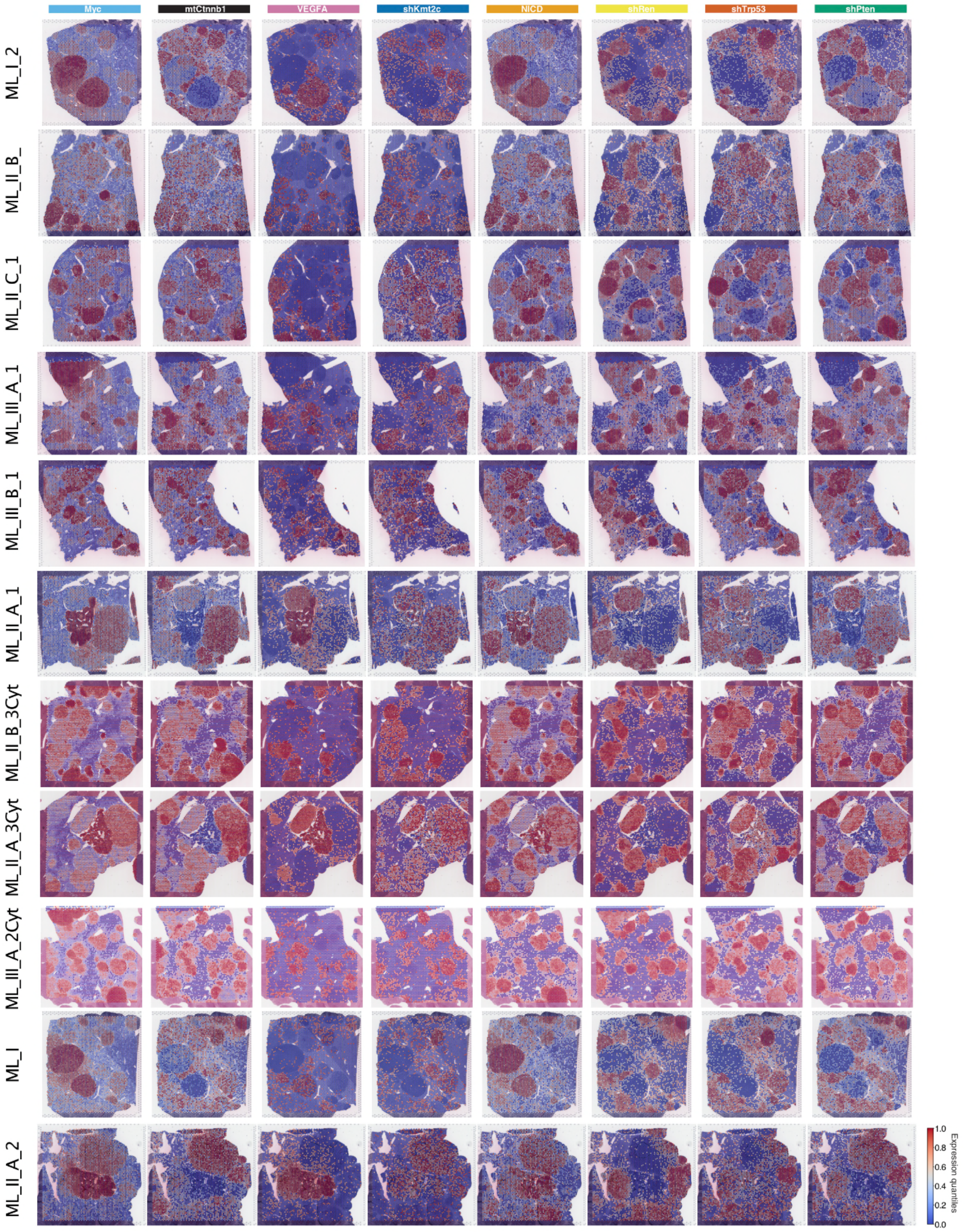


**Extended Data Fig. 7: Spatially-resolved triplet barcode expression enables identification of all 8 perturbations used.**

Spatially resolved expression of triplet barcodes for each of the 8 perturbations across all 11 samples in this study. Gene expression is log1p-transformed and quantile-rescaled, as in Fig. 1c. The quantitative barcode expression for all samples can be explored through the interactive web browser (https://chocolat-g2p.dkfz.de/).

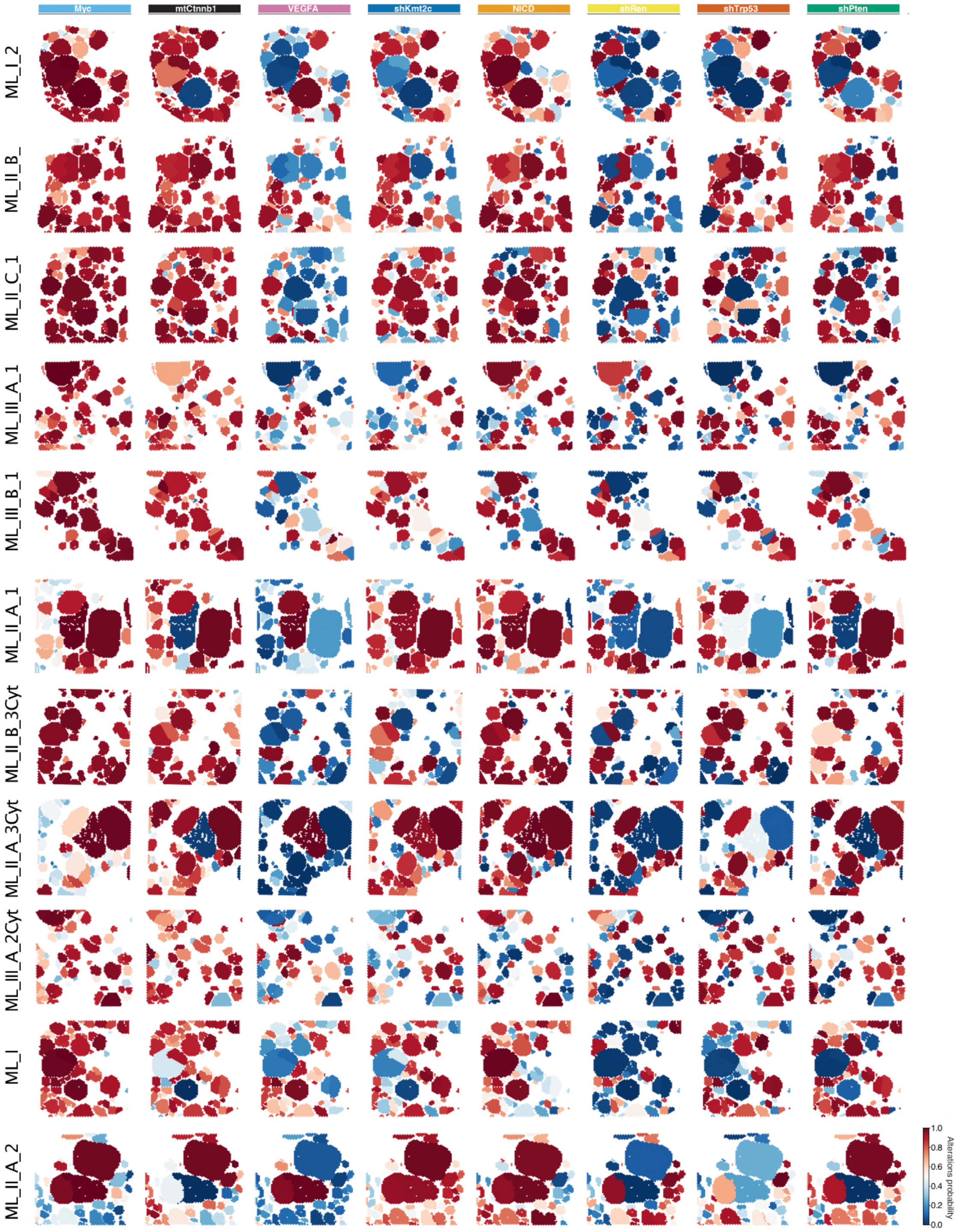


**Extended Data Fig. 8: Spatially-resolved perturbation mapping**

Spatially resolved visualization of the inferred probabilities indicating the presence or absence of each of the 8 perturbations associated with annotated tumor nodules for all 11 samples used in this study (Methods). As in Fig.1d. Perturbation probabilities for all samples can be explored through the interactive web browser (https://chocolat-g2p.dkfz.de/).

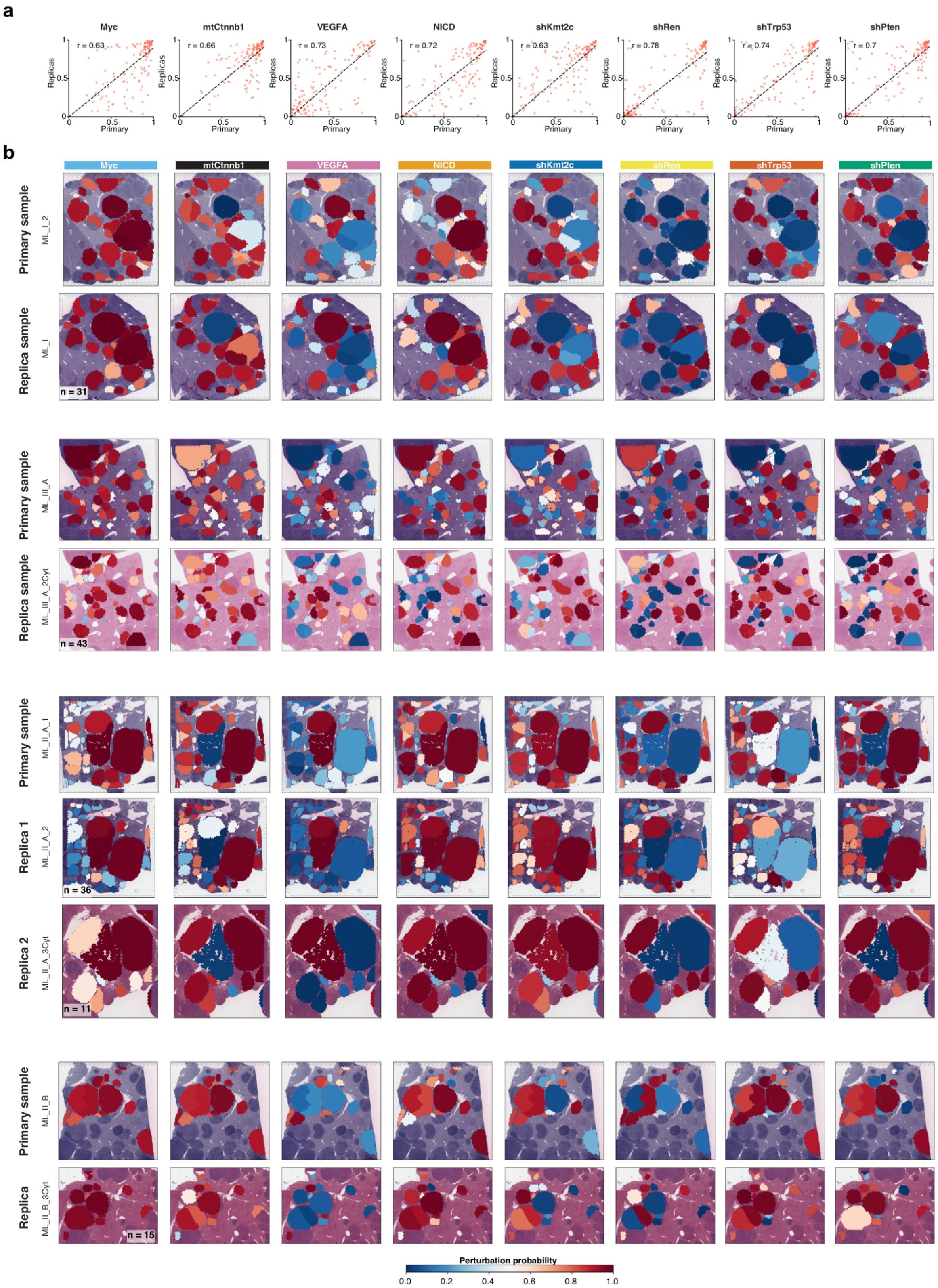


**Extended Data Fig. 9: Quantitative reproducibility of spatial perturbation mapping**

**a, Quantitative reproducibility.** Scatterplots of inferred perturbation probabilities for nodules on the primary section to those on the corresponding replica sections, with Pearson's correlation values displayed. In total, 136 nodule pairs were analyzed.

**b, Matching nodules across samples.** Spatial maps of the inferred probabilities (as in Fig.1d) indicating the presence or absence of each of the 8 perturbations associated with annotated tumor nodules for all samples that have matching ROIs. Matching nodules were manually annotated (Methods). Addition of "_Cyt" in sample name indicates use of 10X CytAssist. Number of matching nodules is indicated.

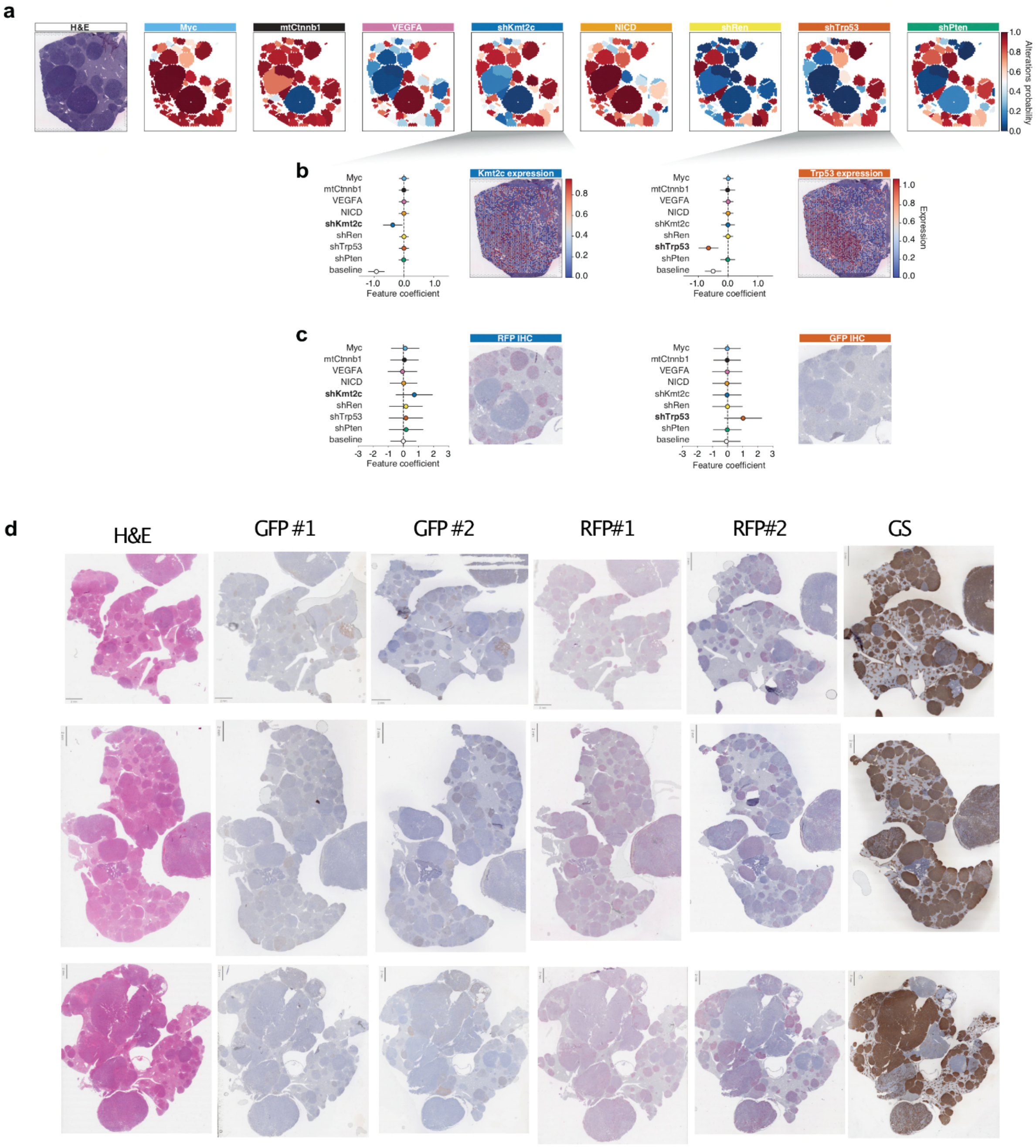


**Extended Data Fig. 10: Validation of inferred perturbation integration**

**a,b,c, Perturbation-phenotype association.** A generalized linear model (GLM) predicts phenotype expression signals based on the estimated probabilities of perturbation presence (Methods). Phenotypes are defined as direct target transcripts associated with perturbations such as shKmt2c-*Kmt2c* and shTrp53-*Trp53* **(b)**. Expression data are log1p-transformed. Note that shTrp53 is linked to a GFP peptide barcode and shKmt2c is linked to a RFP barcode (Extended Data Fig.3). Hence we infer shTrp53-GFP-positive phenotype and shKmt2c-RFP-positive phenotype **(c)**. Representative IHCs for a corresponding ROI on a serial section. Baseline depicts background phenotype marker expression. Error-bars depict 3σ CI. As in Fig. 1e.

**d, H&E and IHC for GFP, RFP, GS.** Three representative FFPE samples were sectioned and stained for H&E (see Extended Data Figure 4). GFP and RFP IHC staining was performed on two individual serial sections. GS IHC staining was performed on serial sections. GFP and RFP were embedded in perturbation plasmids as orthogonal barcodes (see Methods and Extended Data Fig. 3). GS is a well-known marker for liver WNT/mtCtnnb1-signaling activity (see Fig.1e).

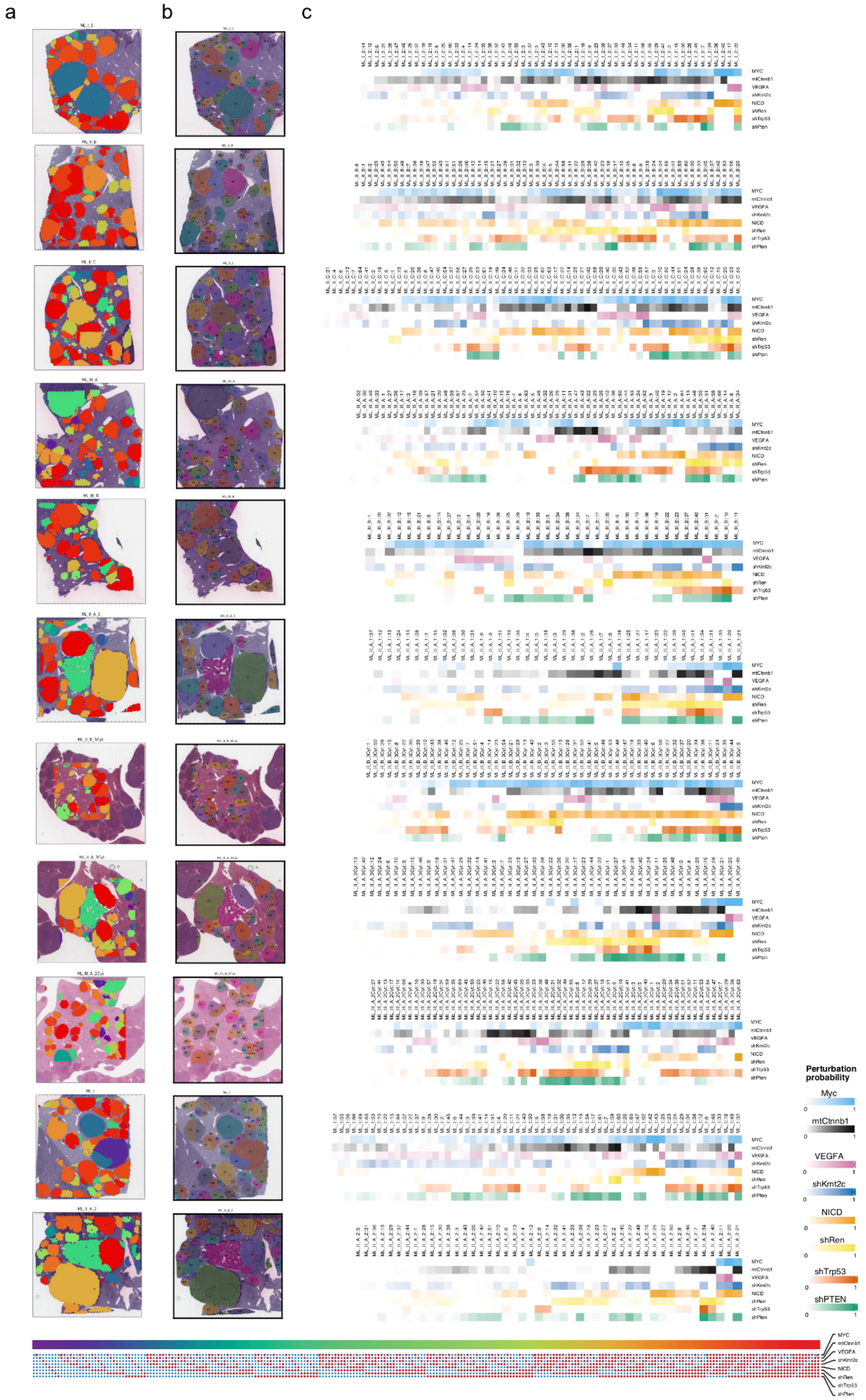


**Extended Data Fig. 11: CHOCOLAT-G2P enables spatial genotype mapping**

**a, Converting barcode signals to genotype maps.** $2^8$ powerset embedding of spatially-mapped perturbations for all 11 samples used in this study, encompassing 622 nodules. Each of the 256 genotypes are color-coded. As in Fig.2a.

**b, Spatially-resolved nodule annotation.** For all 11 samples used in this study. As in Fig.1b.

**c, Tumor genotypes.** Scaled (p^10) estimated plasmid probabilities per nodule. As in Fig. 3.

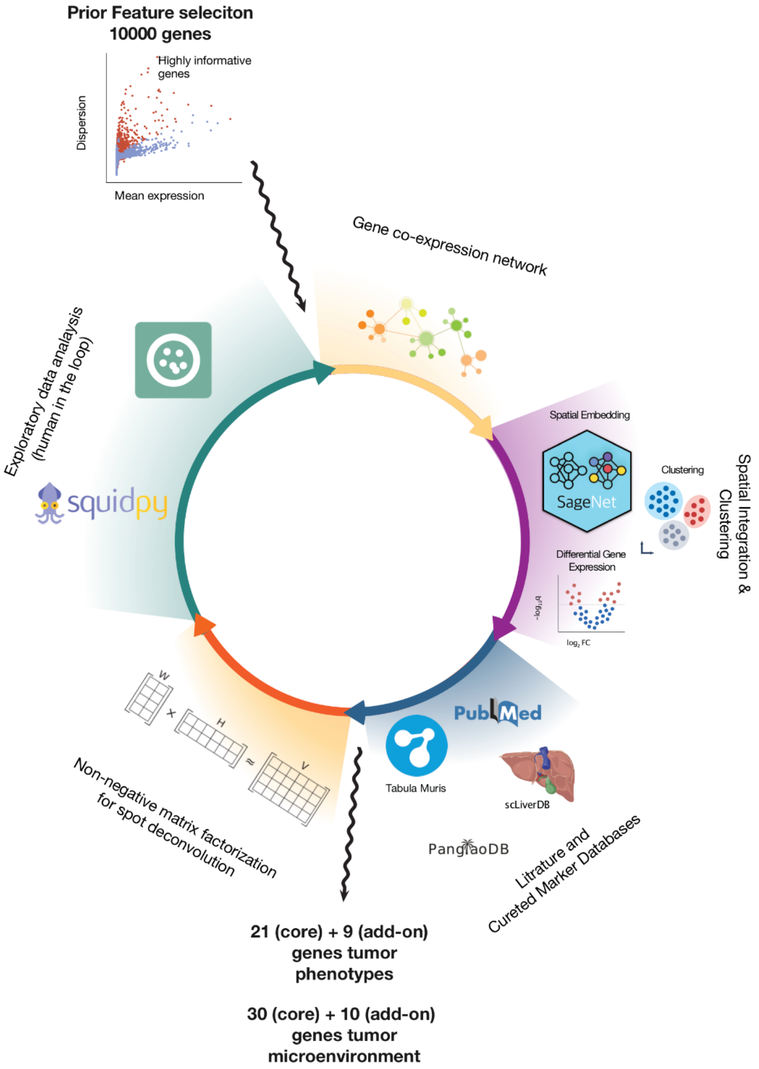


**Extended Data Figure 12: Isolating concise and informative sets of transcripts to characterize cell states.**

Beginning with a pool of 19,510 expressed genes across all ST samples, we identified the top 10,000 highly variable genes per sample, resulting in an overlap of 7,251 highly variable genes across 11 samples. Subsequently, we employed iterative processes involving hybrid data- and knowledge-driven approaches, along with manual verification of consistent gene expression patterns across samples and phenotypes. Through this process, we curated 21 “core marker” genes and 9 additional genes associated with tumor-intrinsic phenotypes, as well as 30 “core marker” genes and 10 additional genes linked to TME phenotypes.

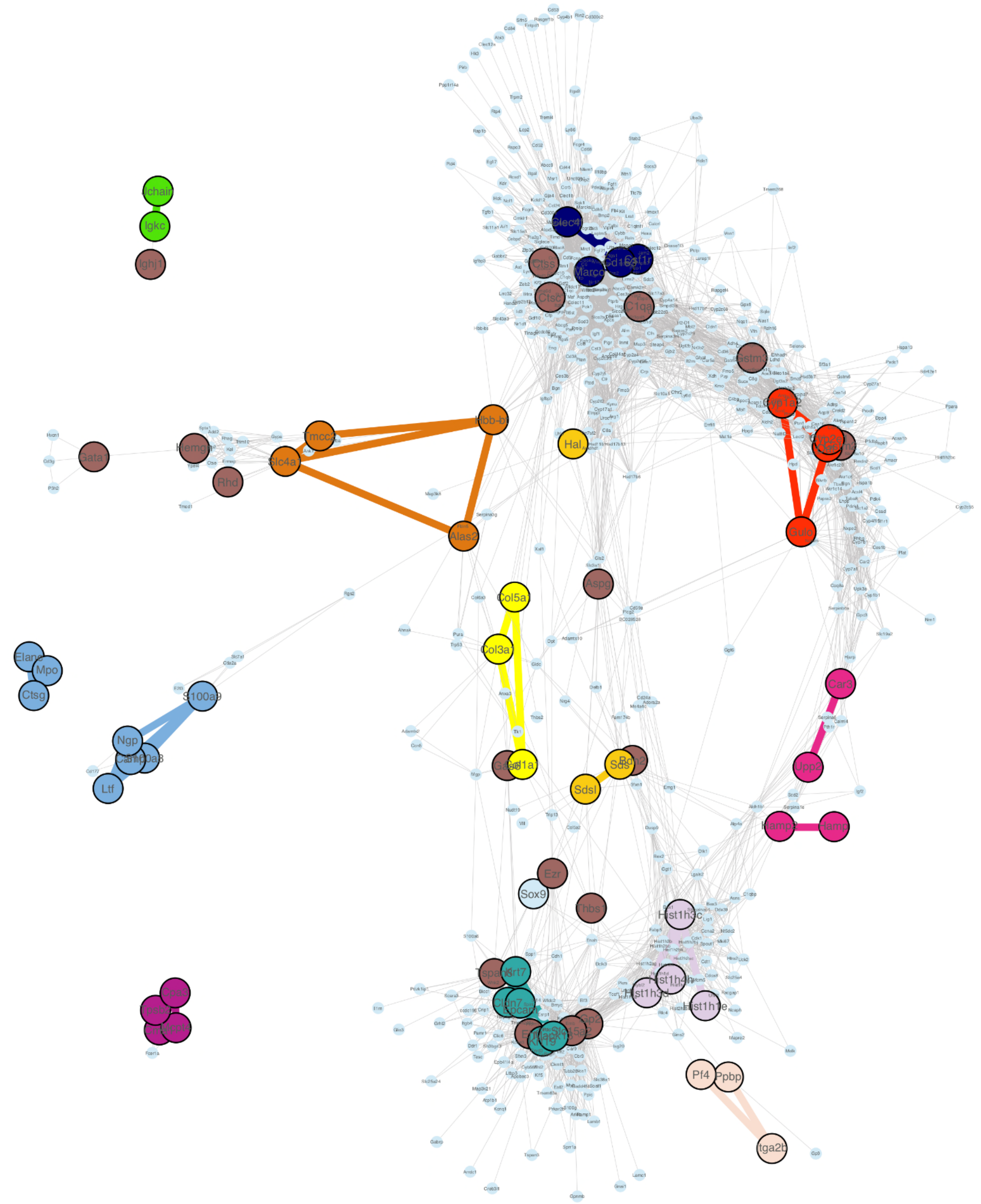


**Extended Data Fig. 13: Transcript co-expression indicates phenotypic relationships and supports marker selection.**

Following the selection of phenotypic “core markers” and associated transcripts, we identified an intersection comprising 500 highly variable genes across all samples. Subsequently, we conducted a Gaussian graphical model analysis (Methods). Nodes correspond to genes, and edges signify significant statistical relationships between pairs of genes. The node colors for “core markers” are indicative of their associated phenotype (see Fig. 3), while associated genes are depicted in brown.

Edges connecting “core marker” genes of the same phenotype are emphasized. Isolated nodes lacking any assigned edges are excluded from the graph.

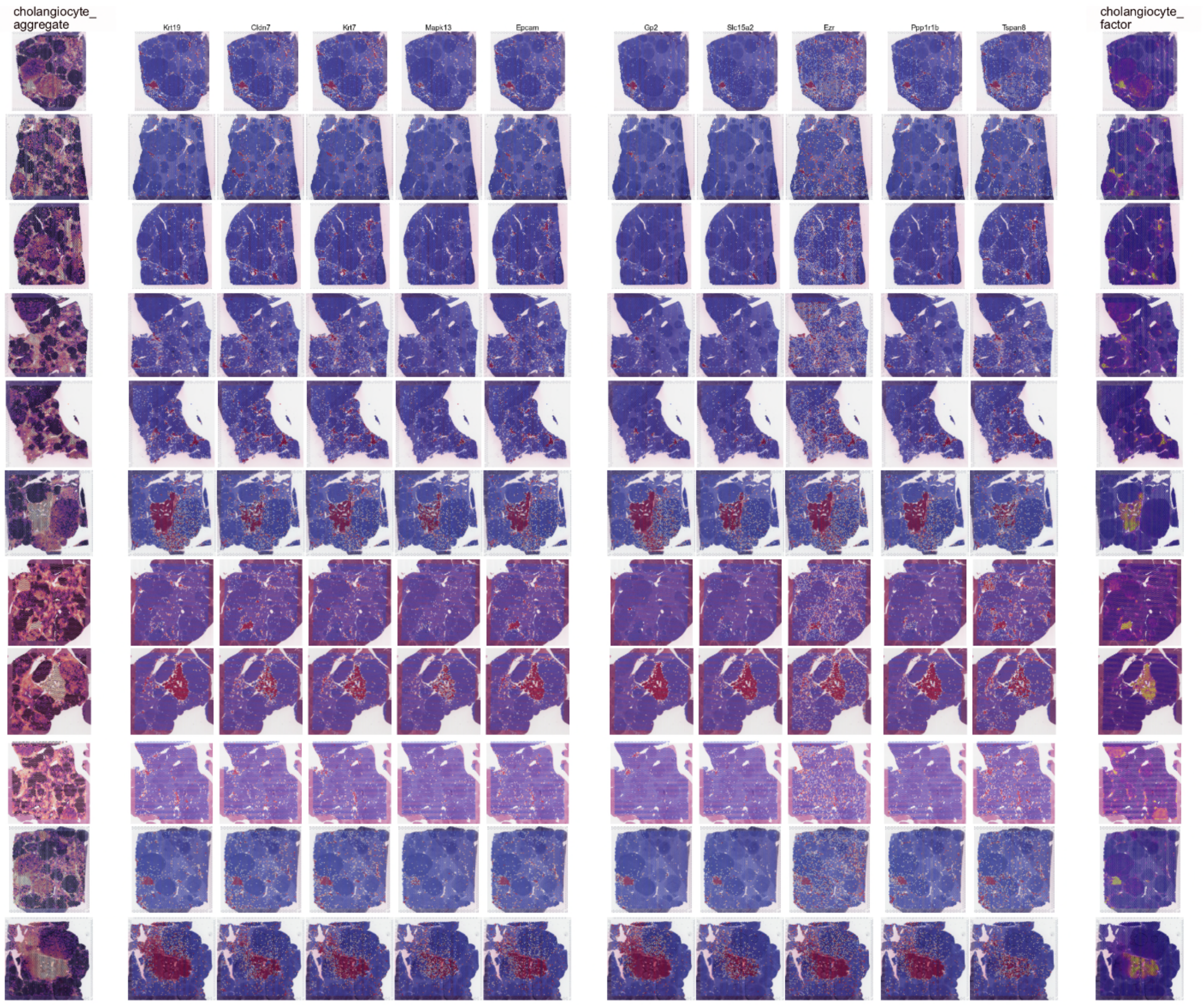


**Extended Data Fig. 14 a, cholangiocyte-like phenotype**

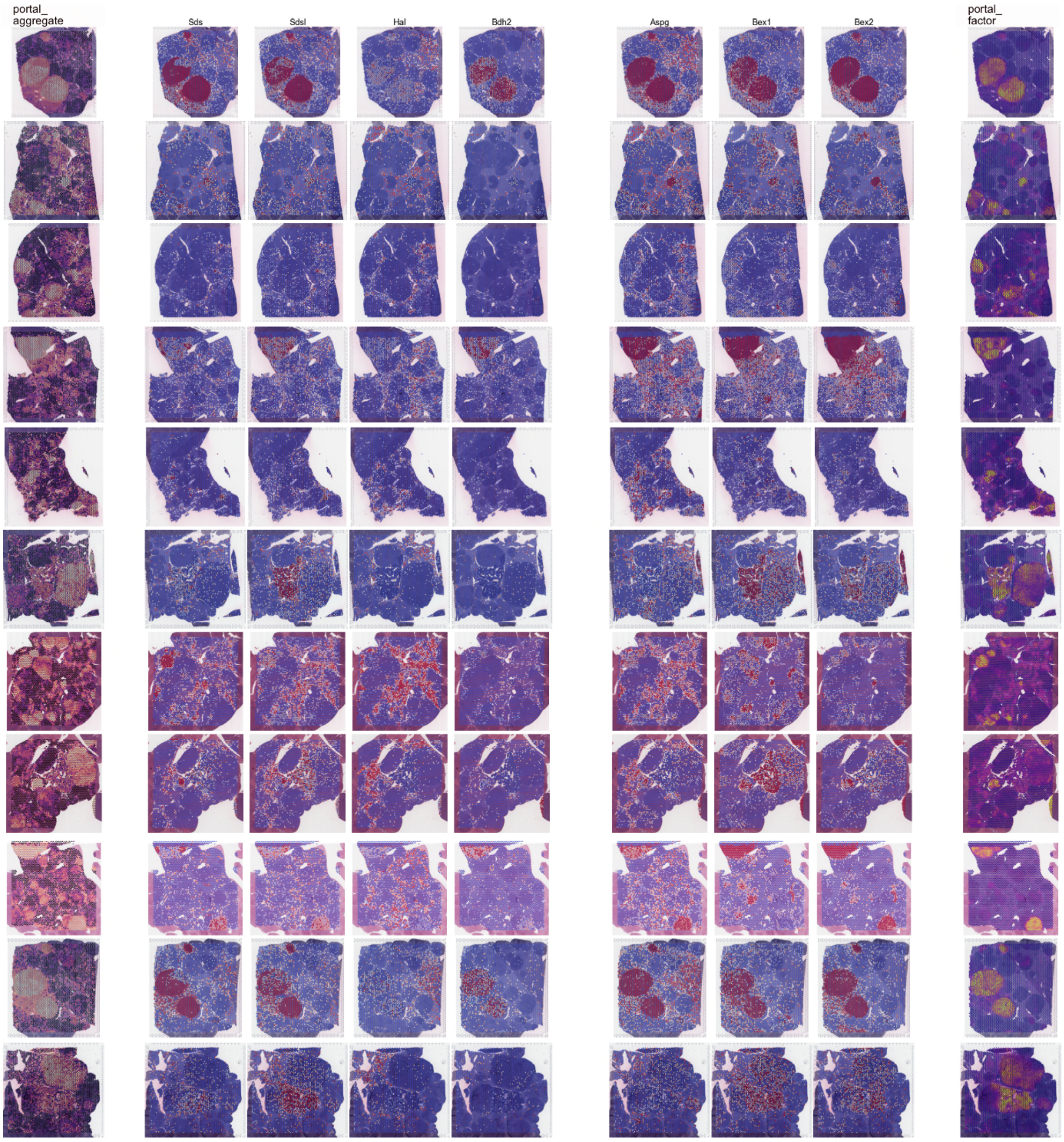


**Extended Data Fig. 14 b, portal-like phenotype**

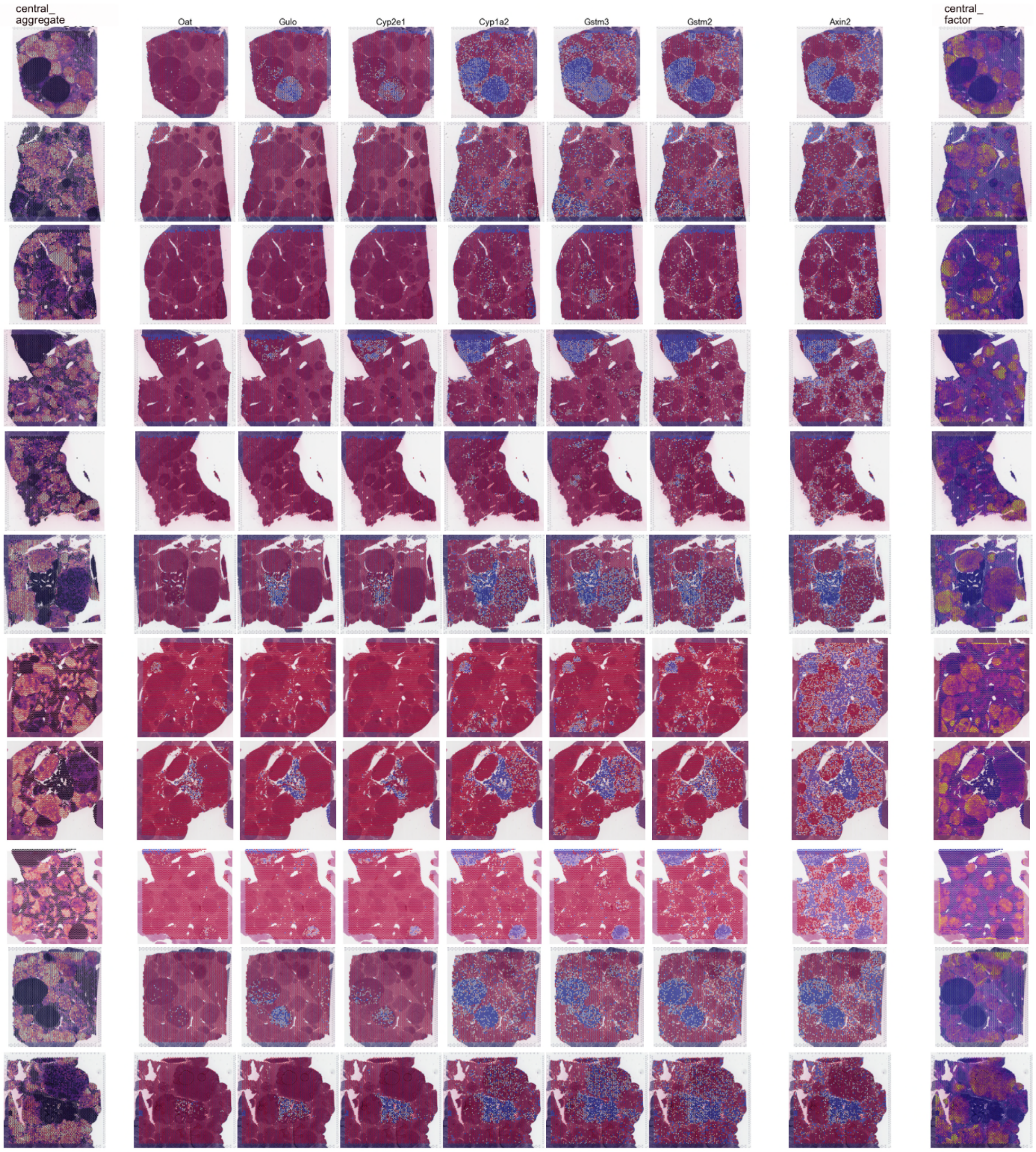


**Extended Data Fig. 14 c, central-like phenotype**

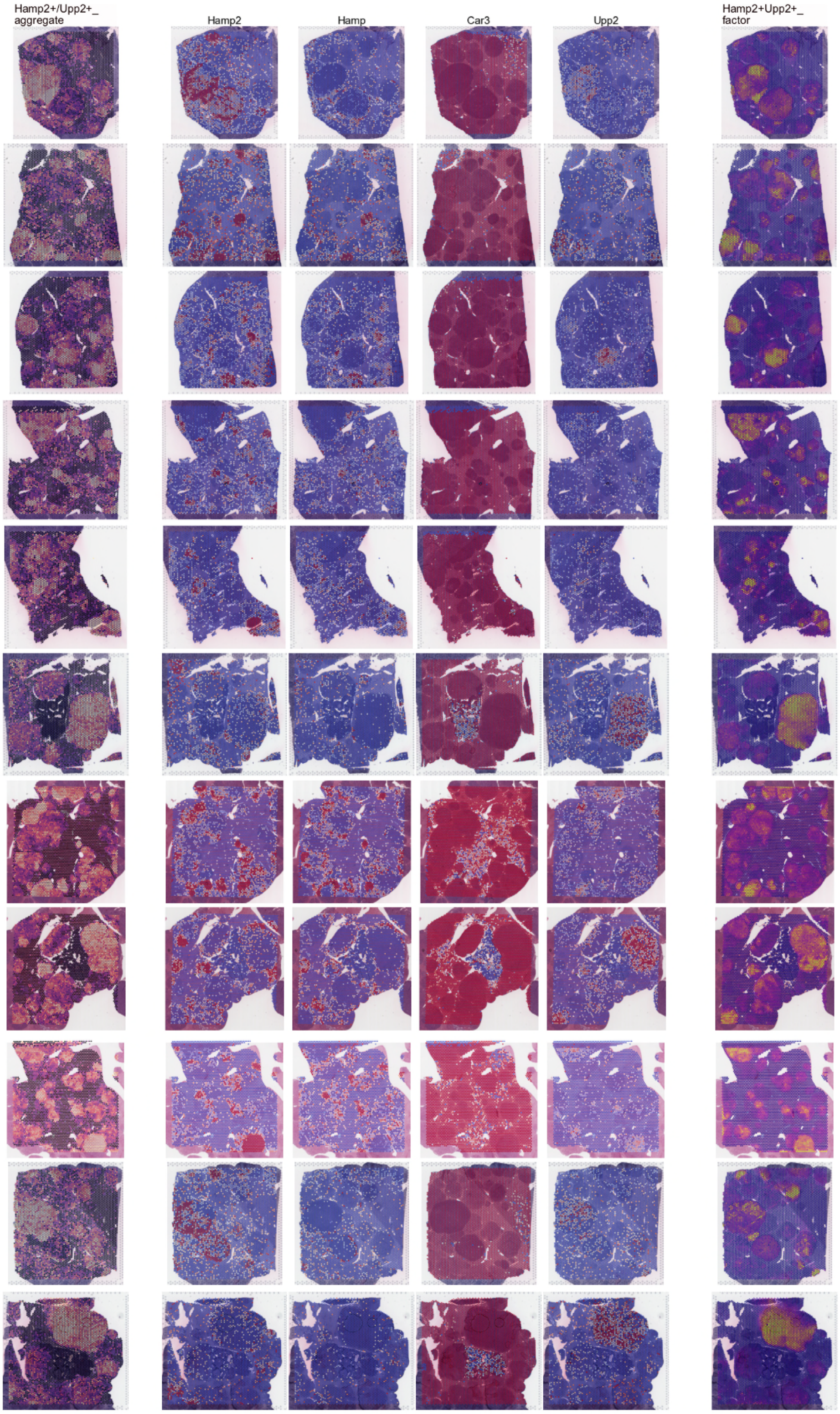


**Extended Data Fig. 14 d, hamp2+/upp2+ phenotype**

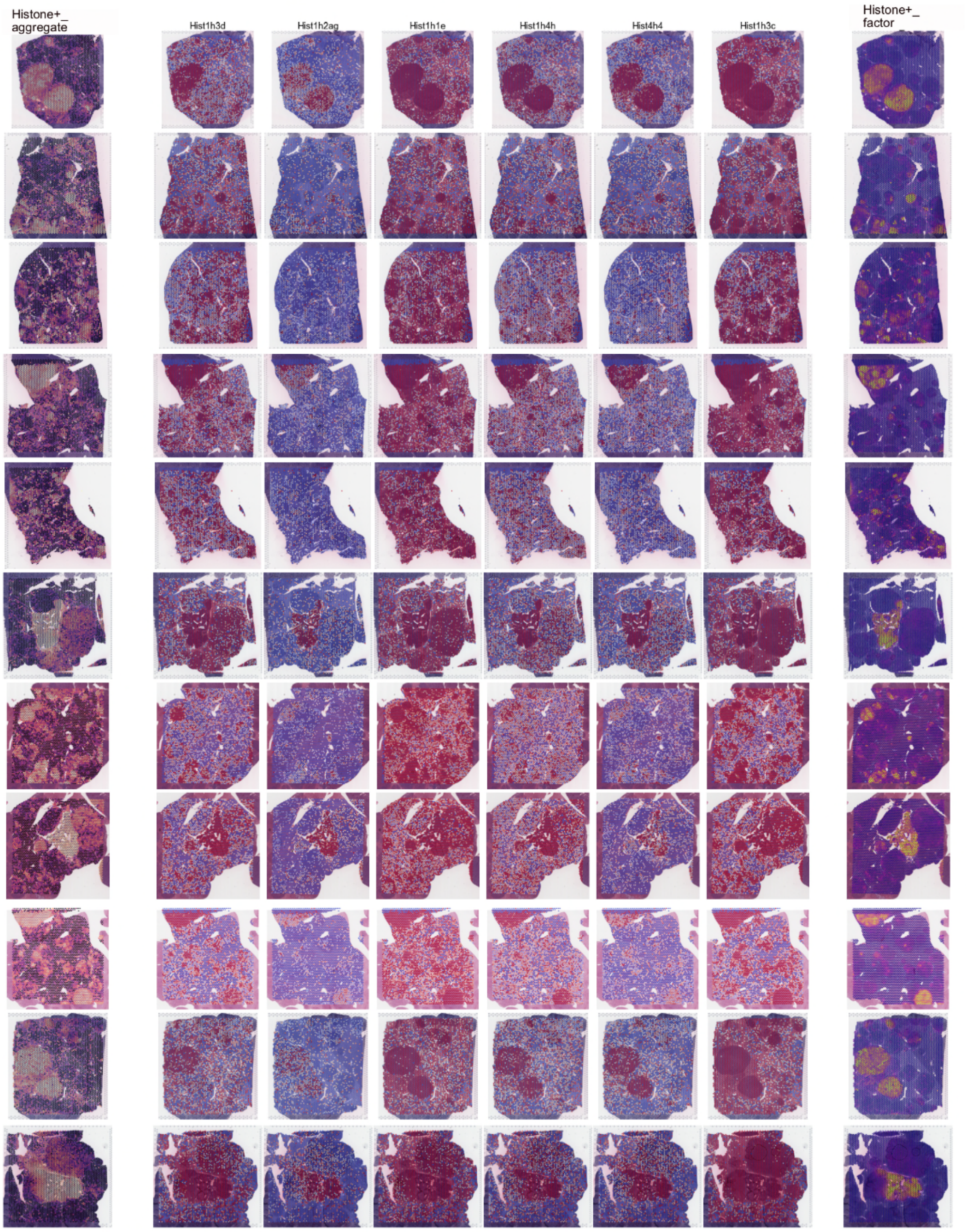


**Extended Data Fig. 14 e, histone+ phenotype**

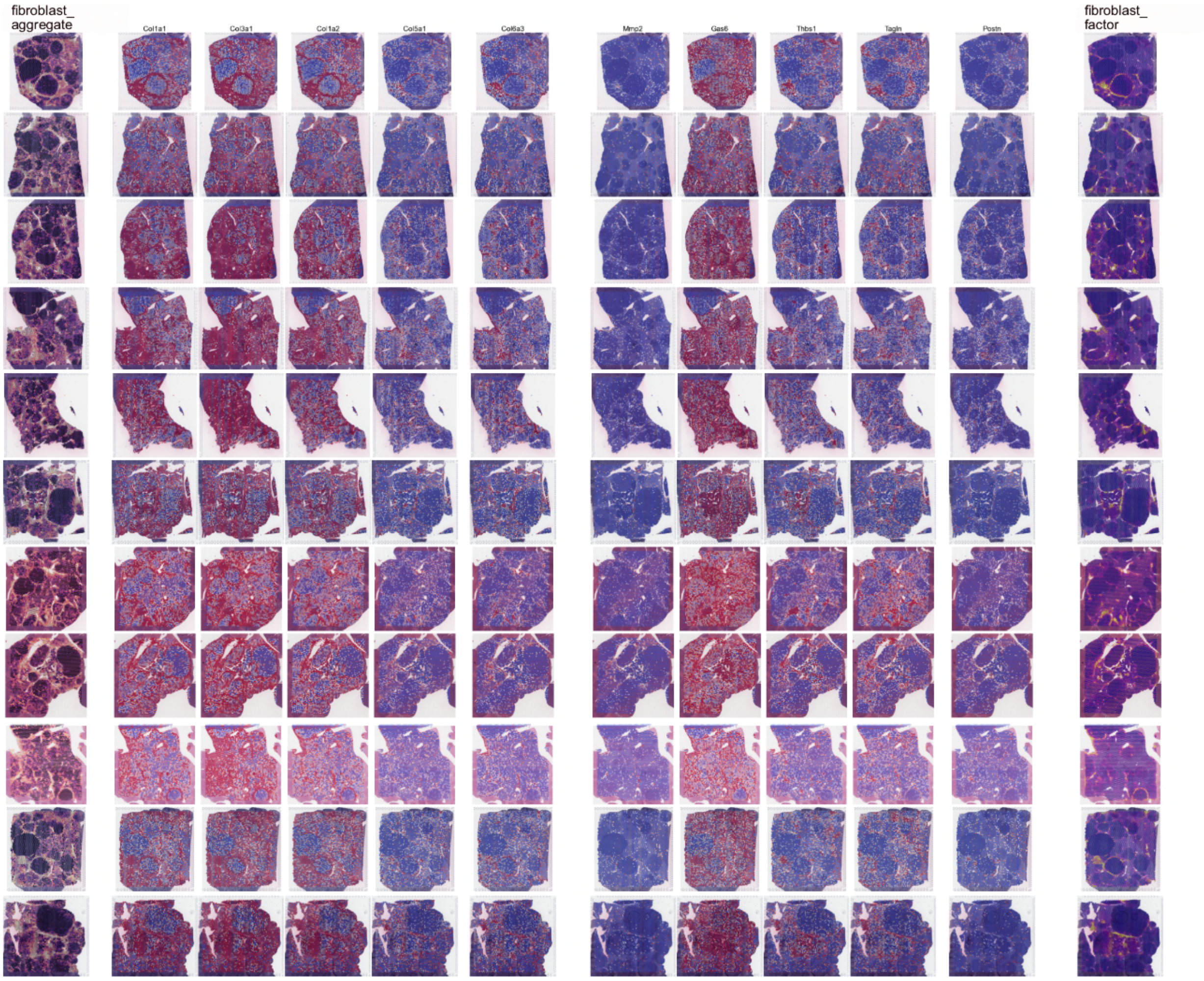


**Extended Data Fig. 14 f, fibroblast-like phenotype**

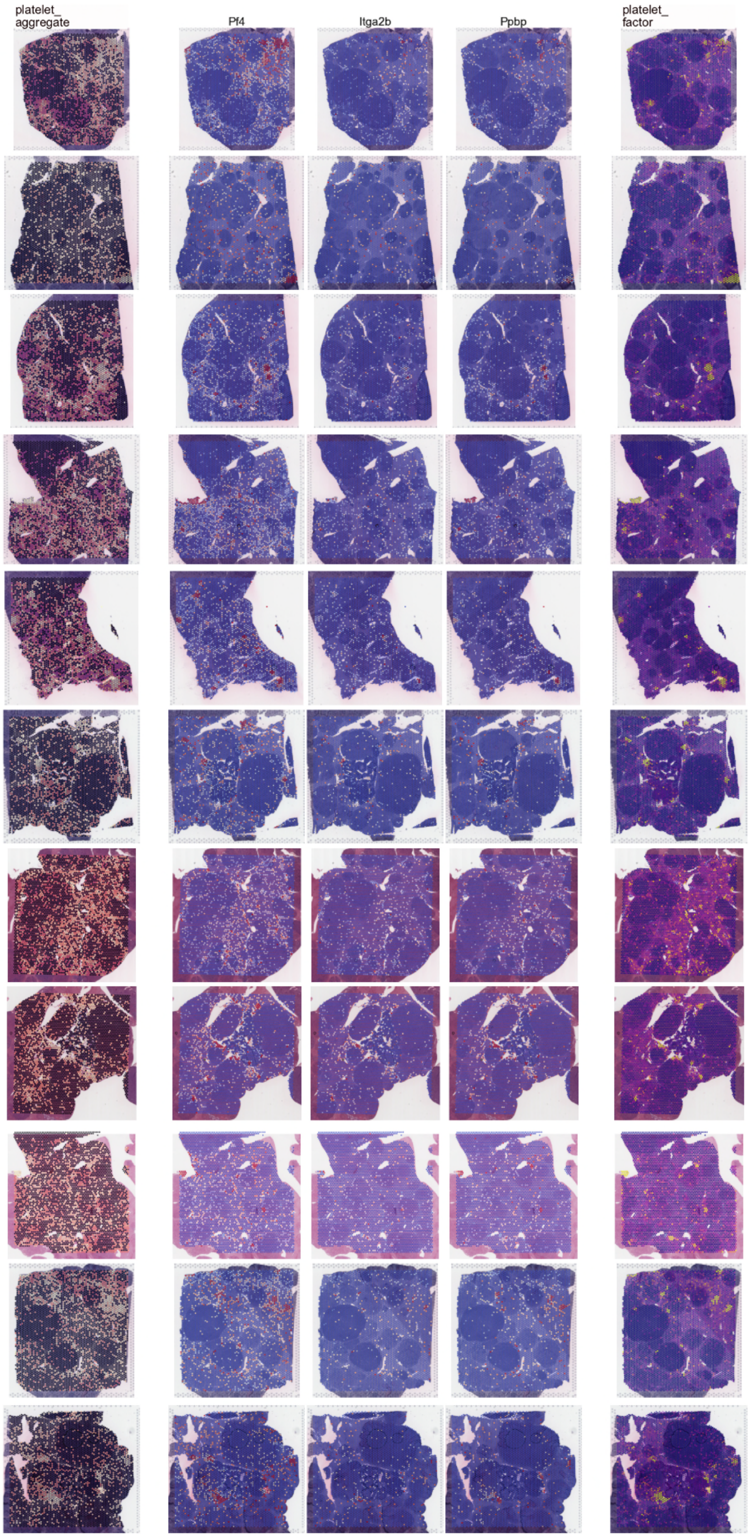


**Extended Data Fig. 14 g, platelet-like phenotype**

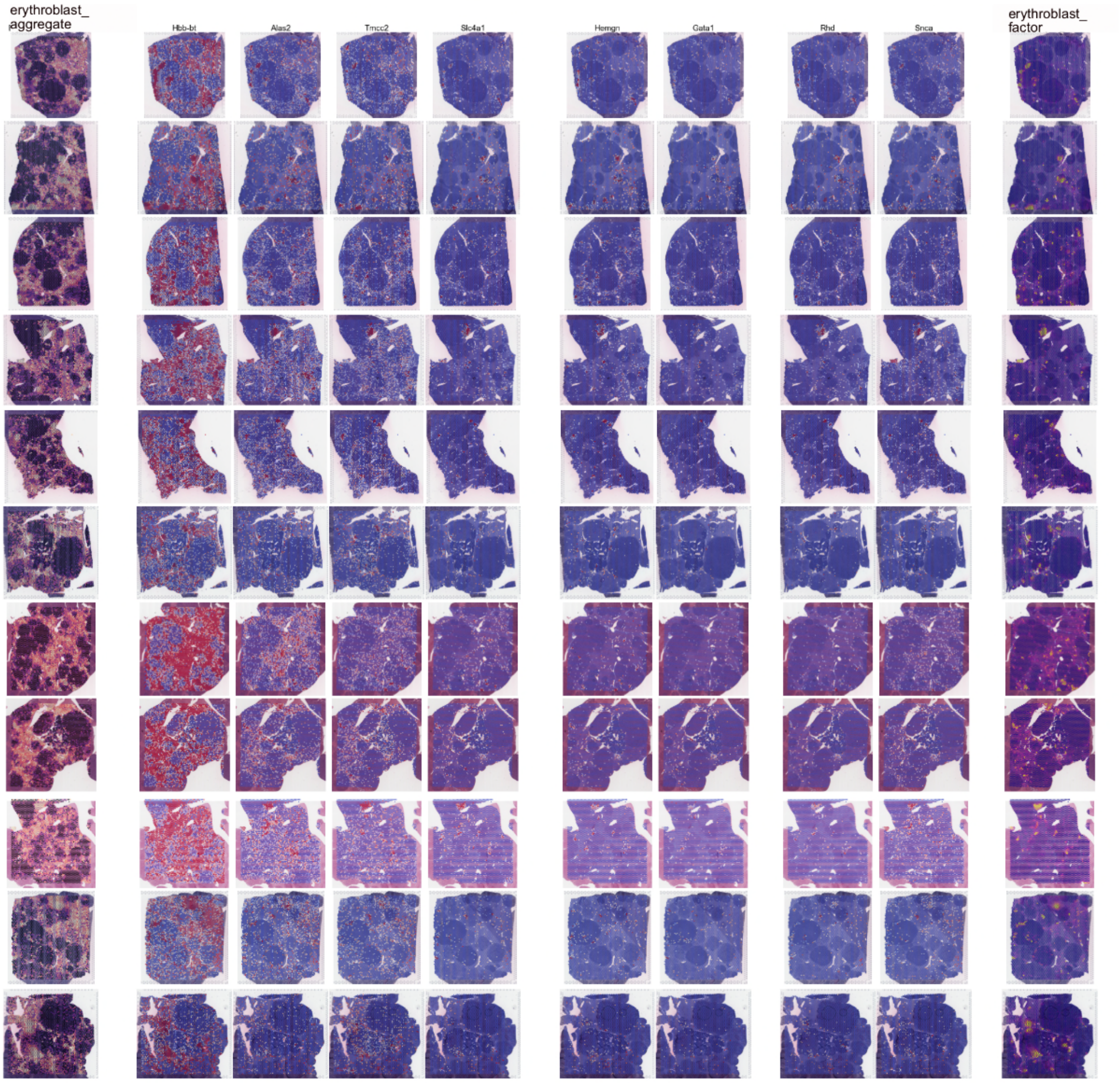


**Extended Data Fig. 14 h,  erythroblast-like phenotype**

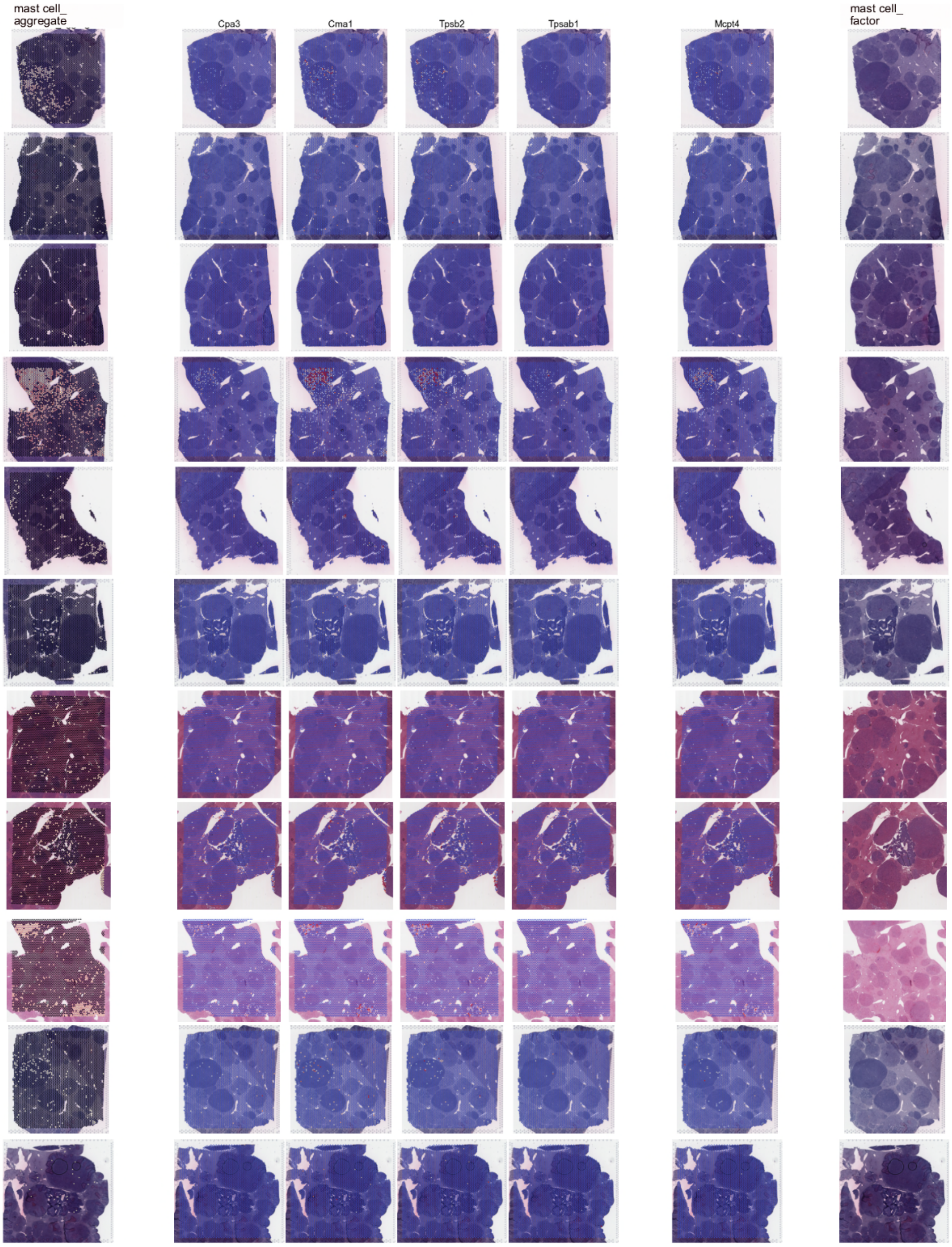


**Extended Data Fig. 14 i,  mast cells-like phenotype**

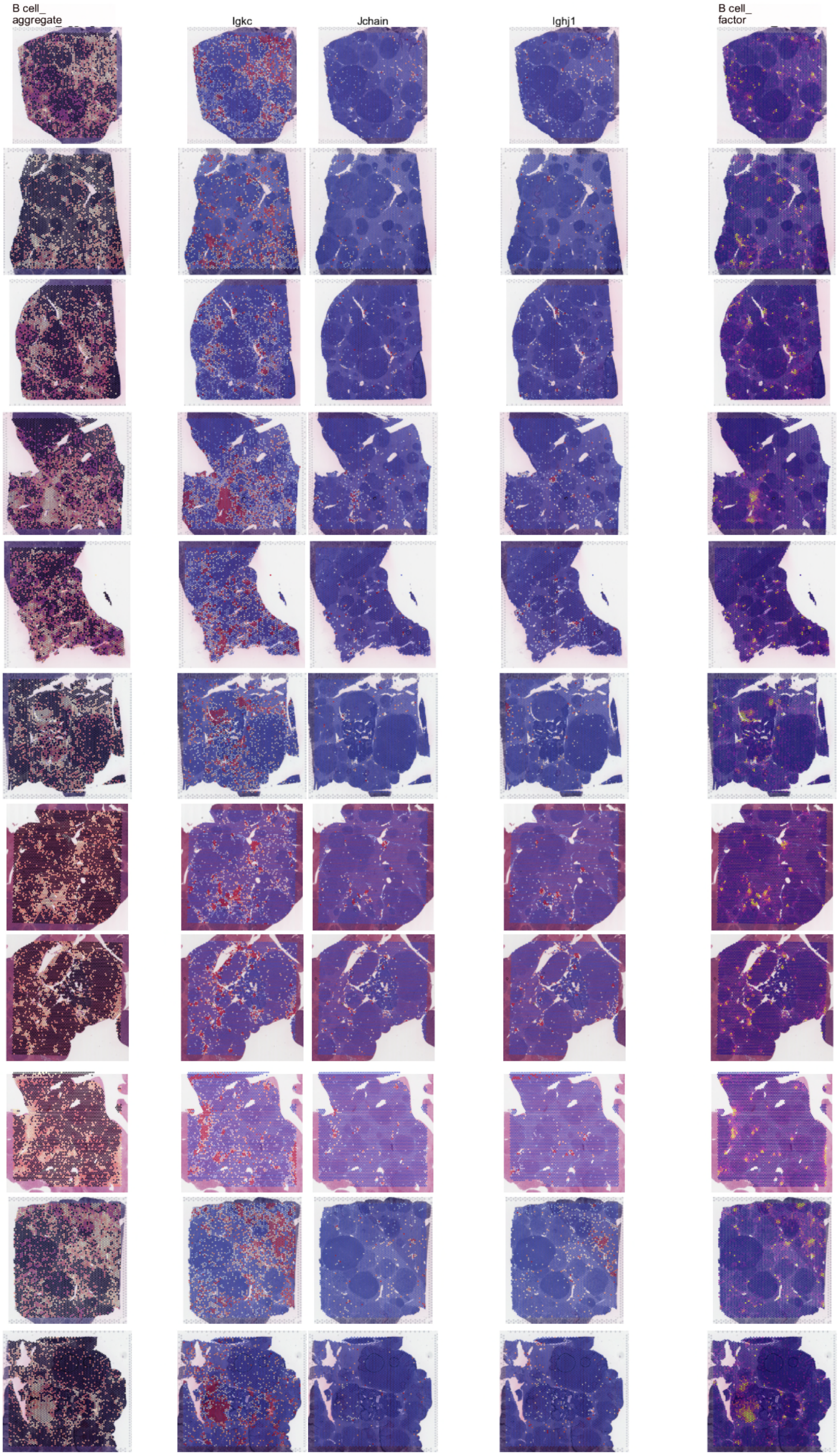


**Extended Data Fig. 14 j, B-cell-like phenotype**

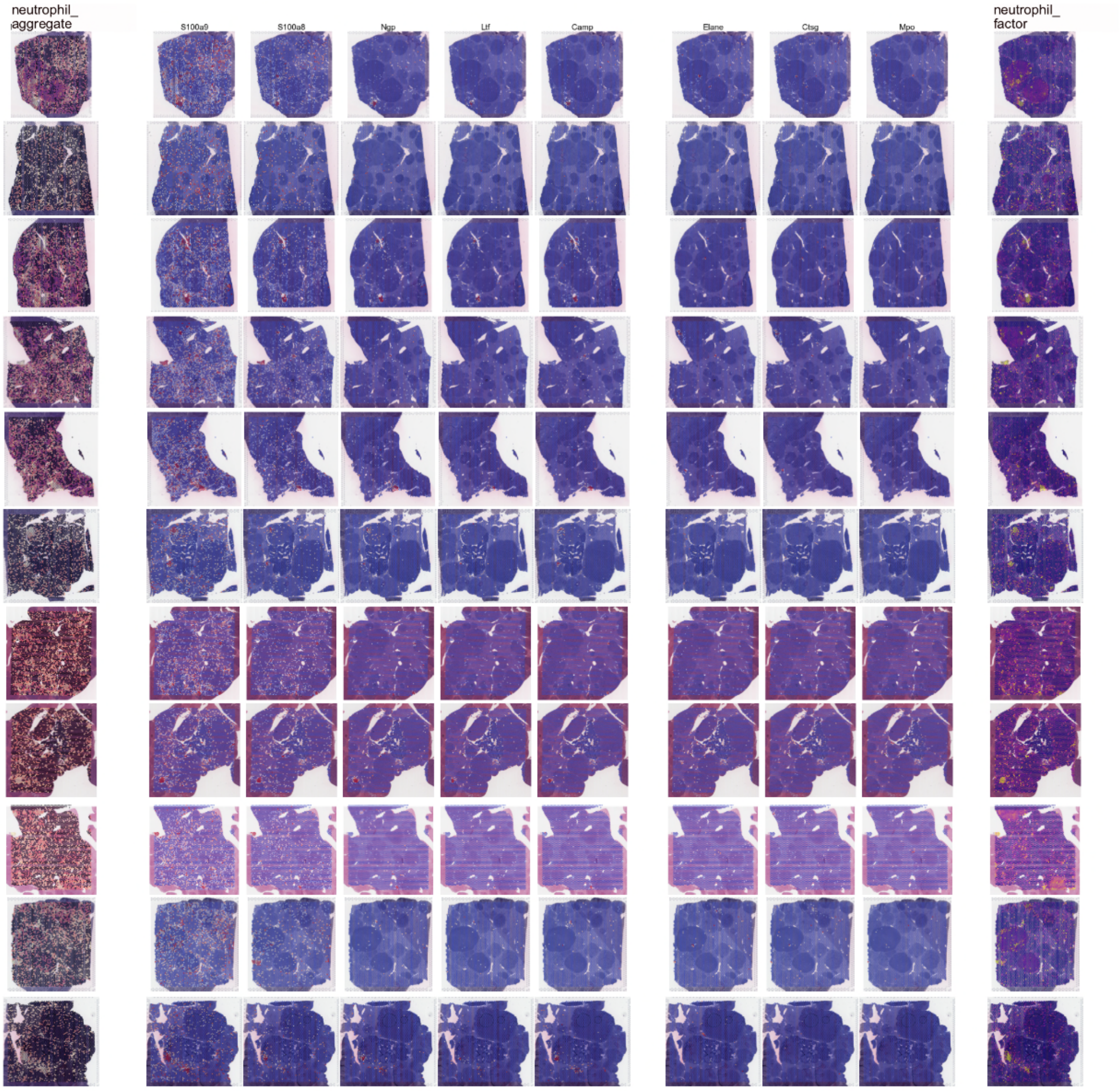


**Extended Data Fig. 14 k, neutrophil-like phenotype**

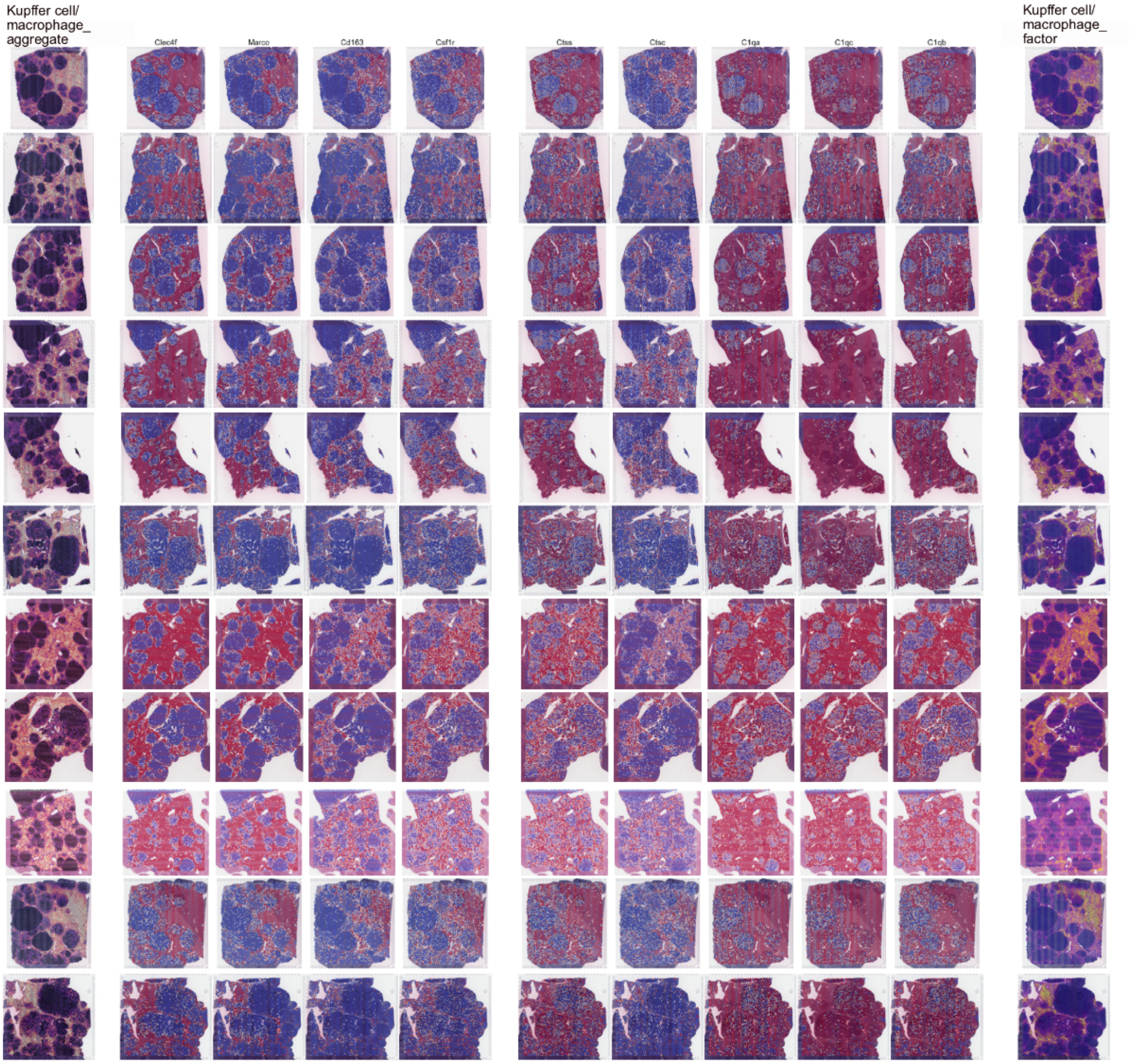


**Extended Data Fig. 14 l, kupffer cell/macrophage-like phenotype**

**Extended Data Fig. 14: Aggregated phenotypes, expression of individual phenotype-associated transcripts, and factor loadings per phenotype of interest.**

**a,** cholangiocyte-like, **b,** portal-like , **c,** central-like, **d,** hamp2+/upp2+, **e,** histone+ **f,** fibroblast-like **g,** platelet-like **h,** erythroblast-like **i,** mast cells-like **j,** B-cells-like **k,** neutrophil-like **l,** kupffer cell/macrophage-like. Left-most column: Aggregated transcript values per each phenotype. As presented in Fig. 3a. Right-most column: Estimated factor loadings per spot (Methods). Columns in between: Scaled expression of single "core markers", as well as associated transcripts. As presented in Fig. 3. Data are depicted for all 11 samples used in this study. See interactive web browser (https://chocolat-g2p.dkfz.de/).

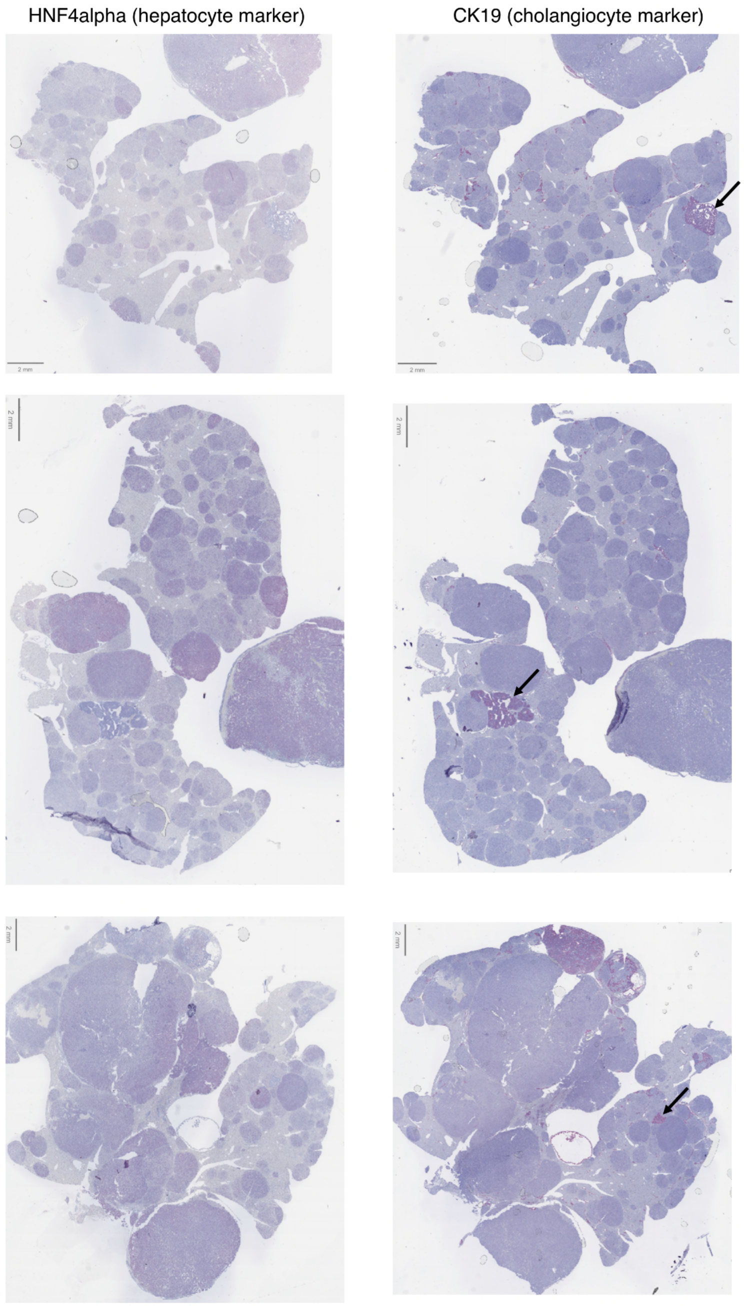


**Extended Data Fig. 15: Immunohistochemistry-based identification of cholangiocytes/cholangiocarcinoma**

Immunohistochemistry for HNF4a (hepatocyte marker) and CK19 (cholangiocyte marker). Arrows indicate prominent cholangiocarcinoma-nodules as an example.

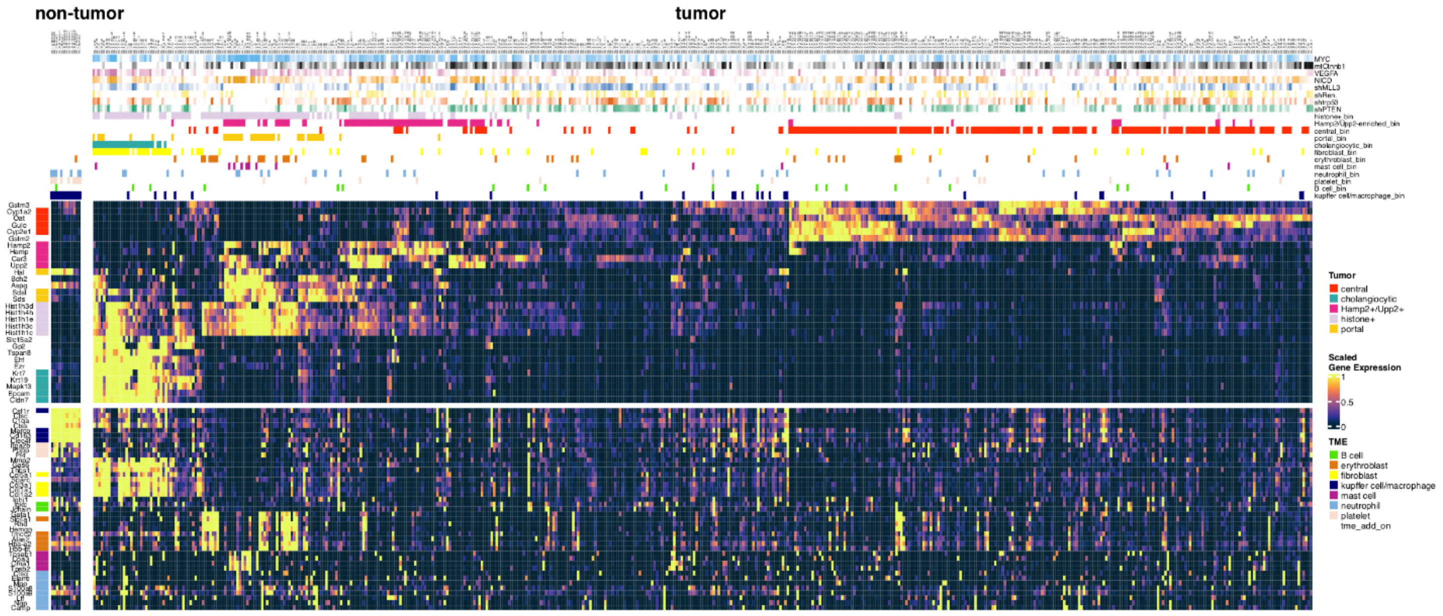


**Extended Data Fig. 16: Overview of genotypes and phenotype-binarization for all nodules across 11 samples.**

As in Fig. 3b and 3d. Top panel: Scaled plasmid probabilities and per-nodule phenotype binarization as used in Fig. 3e (Methods). n=622 total nodules.

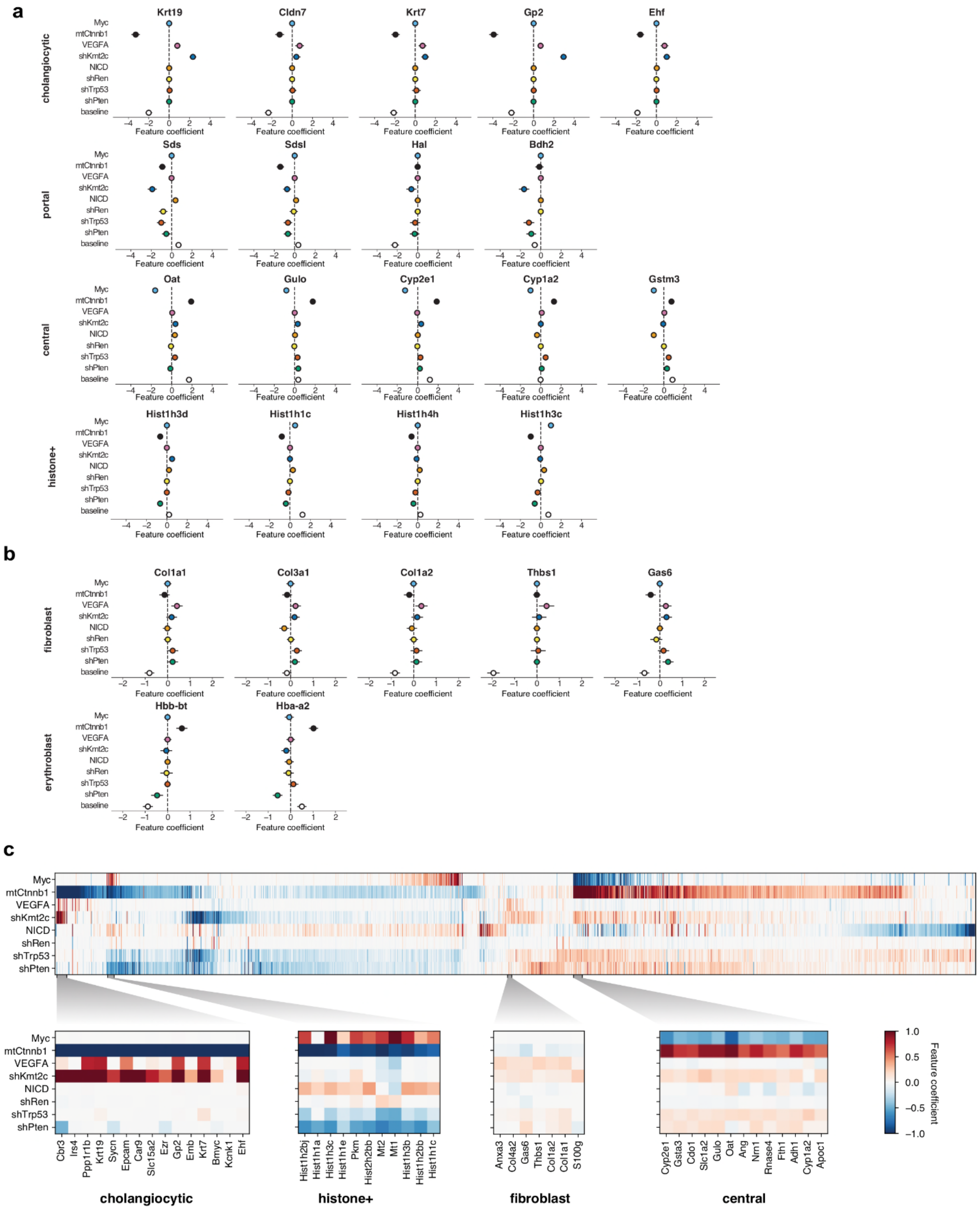


**Extended Data Fig. 17:: Genotype-phenotype relations as evidenced by GLM**

**a, Tumor-intrinsic genotype-phenotype relations.** A generalized linear model (GLM) predicts gene expression signals at each 10X Visium spot, using estimated probabilities of perturbation presence (Methods). Feature coefficients, shown as mean and 3σ confidence intervals, indicate associations between gene expression and perturbations for representative transcripts of four tumor-intrinsic phenotypes.

**b, TME-related genotype-phenotype relations.** As in **(a)** for representative transcripts of two exemplary tumor microenvironment (TME) phenotypes.

**c, Unbiased GLM-inferred genotype-phenotype associations.** Top: heatmap of 1283 genes with at least one significant (3σ) feature weight, ordered by 1D UMAP embedding (see Supplementary Table S2). Bottom: Detailed views of four representative clusters linked to marker genes of known phenotypic groups.

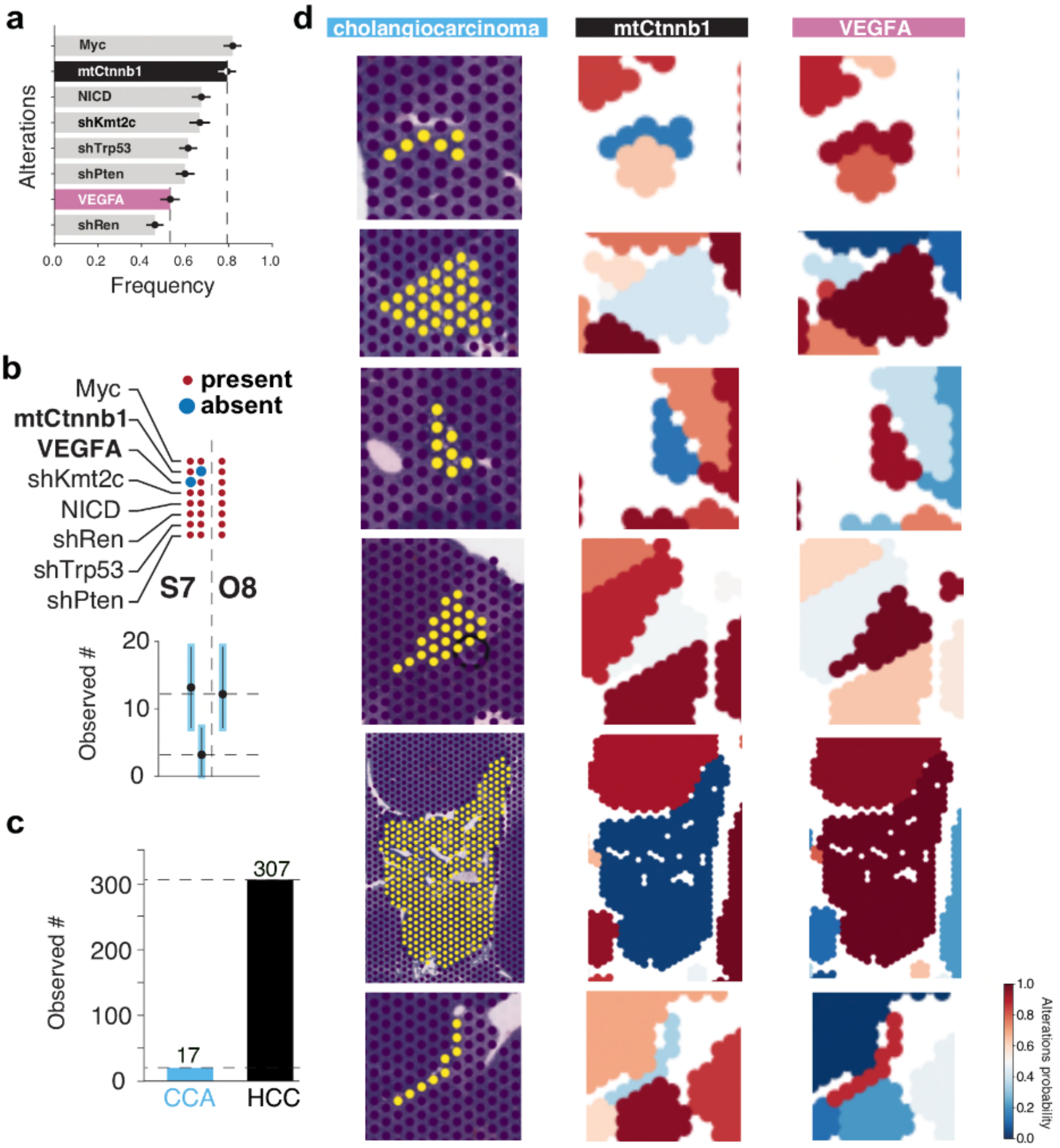


**Extended Data Fig 18: CHOCOLAT-G2P identifies VEGFA and mtCtnnb1 to inversely influence cholangiocarcinoma development.**

**a, Cancer-driving properties.** Frequencies as in Fig. 2d, mtCtnnb1 and VEGFA are highlighted.

**b, Raw tumor occurrence** as in Fig. 2b. Upper panel: perturbation combinations (septets (S7) versus octett (O8)) focusing on absence of mtCtnnb1 or VEGFA; lower panel: raw observed genotypically-defined tumor occurrence (+/- CI) aligned along the 2^8 powerset embedding.

**c, Cancer subtype distribution.** Number of nodules stratified in cholangiocarcinoma (CCA) and hepatocellular carcinoma (HCC; i.e. including all non-CCA subgroups identified in Fig. 3).

**d, Spatially resolved co-occurence of VEGFA and mutual exclusivity of mtCtnnb1 for the cholangiocarcinoma tumor subtype.** Zoom-in of representative nodules identified as cholangiocarcinoma are depicted. Left panel: nodules identified as cholangiocarcinoma, middle panel: VEGFA perturbation probability as in Fig.1d; right panel: mtCtnnb1 perturbation probability as in Fig.1d.

### ONLINE METHODS:

#### Animal experiments

Group size was determined on the basis of our experience with previous experiments[1,2]. For hydrodynamic tail vein injections, 8 week old female C57Bl/6 animals were purchased from Envigo. 5µg DNA of each of a total of 8 perturbation plasmids (40µg total DNA) was mixed with 20µg CMV-SB13 Transposase (1:2 ratio) prepared in a sterile 0.9% Sodium chloride (NaCl) solution and injected into the lateral tail vein with a total volume corresponding 10% of body weight in 5–7s as described before. 2 animals were used. We labeled this approach **RUBIX** (random unique barcode integration combinatorics), since the perturbation plasmids are equipped with molecular barcodes (Extended Data Fig. 3) and the plasmid mixtures injected allow for all possible combinations to become integrated in the genome of hepatocytes (Fig. 1a). All animals were monitored twice weekly and animal experiments were performed in compliance with all relevant ethical regulations determined in the animal permit. When tumors were palpable (10 weeks), animals were euthanized and livers harvested (Extended Data Fig. 4). As a control group, 2 animals received injection of 40µg pT3-EF1-shRen and 20µg CMV-SB13 Transposase (1:2 ratio) prepared in a sterile 0.9% Sodium chloride (NaCl). For fixation, livers were incubated in 4% paraformaldehyde for 48 hours. Sample processing is illustrated in Extended Data Fig. 4. All animal experiments were approved by the regional board Karlsruhe, Germany.

#### 10X Visium for FFPE Spatial Transcriptomics

Spatial Transcriptomics was performed using the manual 10X Visium workflow for samples embedded in paraffin blocks or the 10X Visium CytAssist workflow for samples already placed on glass slides and stained with H&E (Extended Data Fig. 4.). Both workflows were carried out according to the manufacturer protocol (CytAssist, CG000495, RevC; manual Visium, CG000407, RevD). In short, 5 µm slices were cut from FFPE blocks using a microtome and floated onto a water bath at 42°C until all wrinkles were resolved. For the manual Visium workflow the slice was then placed inside the capture frame of the spatial transcriptomcis slide (M.R). Slices used for the CytAssist workflow were placed on frosted glass slides. Deparaffinization and staining

of the slides was similar between both workflows. After drying the slide, paraffin was removed by a 2 hour incubation at 60°C and a subsequent incubation in xylol. Rehydration was done by sequential washes with decreasing Ethanol concentrations. After rehydration the tissue was stained with hematoxylin and eosin and imaged with a Leica Aperio AT2 at 40X magnification. Following imaging the slides were de-stained by incubation in 0.1N HCl and formalin crosslinks were removed by incubation with TE buffer pH 9.0 for one hour at 70°C for the manual workflow and incubation with decrosslinking buffer for one hour at 95°C for the CytAssist workflow. Afterwards tissue was permeabilized and incubated with RTL-probes for approximately 20 hours at 50°C. Free probes were washed away and the bound probes were ligated followed by washing steps to remove unligated probes. For the manual workflow the probes were released by treating the slices with RNase and a permeabilization enzyme. For the CytAssist workflow the slices were stained with a diluted Eosin solution and placed in the CytAssist together with the Visium spatial transcriptomics slides and incubated for 30 minutes at 37°C with RNase and permeabilization enzyme. For both protocols, the spatial barcode was added to the probes by extending them and the probes were released using a 0.08M potassium hydroxide solution. For the CytAssist workflow a pre-amplification PCR with 8 cycles was done. After clean-up with 1.2X SPRIselect beads, 25% of the product was used as input for the index-PCR. For both protocols a qPCR was done to select the number of cycles for the index PCR. To reduce PCR duplicates and avoid over-amplification, cycle number at a Cq-value of 10% was used for the index PCR. The PCR product was purified using 0.85X SPRIselect beads.
For samples already stained and mounted on a slide, slides were first imaged and then incubated in xylol to remove the coverslip. Sample rehydration was done by sequential washes with decreasing ethanol concentrations. Slides were destained and decrosslinking was performed by incubating with a decrosslinking buffer at 95°C for 1 hour. After decrosslinking, samples were incubated with probes for 20 hours at 50°C. Excess probes were washed away and the probes were ligated. Thereafter, unligated probes were washed away. The samples were stained again with eosin and placed in the 10X CytAssist together with the spatial transcriptomics slides. A mixture of RNase and permeabilization enzyme was added to the spatial transcriptomics slides and the 10X CytAssist was started. After incubation, the spatial transcriptomics slides were removed and the enzymes were washed away. The spatial barcodes were attached to the probes with an extension enzyme. Probes were released using 0.08M

potassium hydroxide solution. Probes were then amplified by 8 PCR cycles. 25% of the purified PCR products were used as input for the index PCR. The cycle number of the index PCR was determined using the cycle number at a Cq-value of 10%.

For all samples the final sample concentration was determined using Agilent Tapestation 4150 with D1000 HS tapes. Sequencing for both protocols was performed on an Illumina NovaSeq6000. Four samples were pooled on one SP flow cell with 100 cycles to aim for a read count of 250M Reads per sample. The fastq files and the alignment was done using spaceranger 2.0.1. A total of 12 ST-datasets were generated (Extended Data Fig. 4). Note that the utility of sample ML-II_B_2Cyt is constrained by tissue detachment of the sample during the processing for 10X Visium CytAssist.

10X Visium for FFPE engages RTL-probes that capture a 50 nt sequence specific to endogenous transcripts (Note that we used Visium Mouse Transcriptome Probe Set v1). We leveraged this strategy to likewise capture molecular barcodes via RTL-probes (Fig. 1a). We hence labeled this approach **PERTURB-CAST** (Perturbation barcode capture spatial transcriptomics).

**Economized molecular barcode selection:**

To enable PERTURB-CAST, we aimed to avoid additional expenses and protocol modifications by redeploying RTL-probes from commercially available 10X Visium reagents (originally designed to detect endogenous transcripts), as barcode identification reagents, provided that the selected transcripts are not expressed in mouse liver. We labeled this strategy **REDPRO-BC** (redeploy probes for barcode capture, Extended Data Fig. 2). To this end, we initially analyzed a publicly available bulk RNA-seq dataset including a total of 128 murine liver samples (GSE137385[3]) to identify transcripts that were generally not detected (FPKM=0 over all samples). Note that this approach can be error-prone due to the initial source data. Consequently, we went on to validate non-expression of selected transcripts (olfactory-, vomeronasal- , taste-receptors) in additional datasets (including bulk-RNAseq from GSE148379[4], information provided in MGI GXD[5] as well as 10X Visium data[6]). The endogenous transcripts associated with the REDPRO-BCs used in this study are illustrated in Extended Data Fig. 3 and respective nucleotide sequences for 10X Visium RTL-probe capture barcodes (reverse complement to RTL-probe sequence provided by 10X Genomics) are listed under the section Molecular cloning. Note that we used Visium

Mouse Transcriptome Probe Set v1. Visium Mouse Transcriptome Probe Set v2 is not compatible with the barcodes employed in this study.

**Molecular cloning**

Transposon plasmids used in this study including overexpression constructs for NICD, mutant human *CTNNB1* (T41A), and human *MYC* and potent shRNA constructs against *Trp53, Pten, Kmt2c* and *Renilla luciferase* were described and validated in animal experiments before[1,4,7]. For ST, note that NICD overexpression can be investigated via *Notch1* expression given that the 10X Visium RTL-probe identifies the exogenous transcript introduced. However, endogenous *Notch1* is likewise identified. VEGFA overexpression plasmid was cloned by insertion of a codon-optimized gene fragment (gBlock, IDT) based on *Vegfa* NCBI Reference Sequence: NP_033531.3 by replacing h*MYC* from a previously validated expression plasmid[1] using NEBuilder HiFi-DNA Assembly according to the manufacturer's protocol. All plasmids were individually modified to express molecular barcodes. Specifically, fluorescent protein-based peptide barcodes (mKate2, mOrange2, mWasabi) were ordered as codon-optimized gene fragments (gBlock, IDT) and cloned into previously validated shRNA expression plasmids to replace GFP[1] using NEBuilder HiFi-DNA Assembly according to the manufacturer's protocol (NEB). Small-peptide barcodes (e.g. AU1, AU5, etc.) as described in[8] were ordered as oligos (Sigma), annealed and cloned using NEBuilder HiFi-DNA Assembly according to the manufacturer's protocol (NEB). Long RNA-barcodes (stretches of at least 650 nts derived from the combination of multiple oligo-miner probe sequences designed against Arabidopsis thaliana Chr1[9]) were ordered as gene fragments (gBlock, IDT) and cloned using NEBuilder HiFi-DNA Assembly according to the manufacturer's protocol (NEB). REDPRO-BC triplet-arrays were ordered as gene fragments (gBlock, IDT) and cloned using NEBuilder HiFi-DNA Assembly according to the manufacturer's protocol (NEB). In brief, each REDPRO-BC triplet-array incorporates one 50 nt sequence derived from olfactory-receptors, one 50 nt sequence derived from taste-receptors, and one 50 nt sequence derived from vomeronasal-receptors (based on 10X Genomics RTL-probe sequences against murine transcripts; Note that we used Visium Mouse Transcriptome Probe Set v1.; see below), separated and flanked by ca. 20 nt spacer sequences to avoid potential steric hindrance during hybridization. Spacer sequences used were derived from T7 and T3 promoters as described in[10] and/or AsCas12a-DR sequences described in[11] and/or 10X Capture sequences cs1 and cs2 as described in[12] and as such provide additional functionality which was not tested in this study. Single 50 nt REDPRO-BCs were ordered as oligos (Sigma), annealed and cloned using NEBuilder HiFi-DNA Assembly according to the manufacturer's protocol (NEB). Note that REDPRO-BC length should enable straightforward en-masse cloning engaging commercially available oligo-pools such as in[11] which was not tested in this study. Peptide barcodes were integrated in frame with respective coding regions. RNA barcodes were integrated in the 3' UTR of coding regions expressed under the control of a polymerase II promoter (EF1) unless otherwise specified. Subsets of perturbation plasmids were equipped with REDPRO-BC arrays either 5' and 3' of the shRNA expression cassette (miRE-based) to account for miRE-processing[13], or were equipped with additional REDPRO-BC arrays driven by a polymerase III promoter (hU6) in reverse orientation to the EF1-driven transcript. Extended Data Fig. 3 illustrates plasmid design. Respective FASTA sequences of plasmids are available upon request. Plasmids were validated by restriction digest and Sanger sequencing (Microsynth).

Selected REDPRO-BC barcode sequences based on Visium Mouse Transcriptome Probe Set v1. were:

**Myc:**

```
Olfr103:  TGGGAGTGAGAGACATACAAGAACCACAGCCCTTTCTCTTTGCTATTTTC
Tas2r102: AACACAAGTGTGAATACCATGAGCAATGACCTTGCAATGTGGACCGAGCT
Vmn1r1:   TAAAAGGCAGTGTCAGTACCTTCACAACACCAGCATTTCCCGCAAAGCAT
Olfr1018: CAGTTCCATGGTTATCAATGTTCTCACCTTGAGTTTGCCCTACTGTGGAC
Tas2r118: TTATTGGCACTGTGTTTGATAAGAAATCTTGGTTCTGGGTCTGCGAAGCT
Vmn1r174: ACTTCAACCAGAGGCCAGAGCAGCAAACACAATTCTCATGCTGATGATCA
Olfr1:    TGGCCAGCATCTTTCTTGTCCTTCCATTTGCACTCATTACCATGTCCTAT
```

**mutant(mt)Ctnnb1:**

```
Olfr1000: GGCACAGTAGGTATGTTCACTGGTCTGATAATTCTGGGGTCCTATGTATG
Tas2r103: TGTCACTAATCACAGGGTTCTTGGTATCATTATTGGACCCAGCTTTATTG
Vmn1r178: GTCTCTTCATGAGTCATTTCAGTAAAGTTTTTGCTGCAGGATTCCCCACT
```

Olfr1019: TGCTTGGTCCTAATGCTGGGCTCTTACTTCGCTGGCCTAGTGAGTTTAGT

Tas2r119: GATATCCAGGTTGGTGCCATGGCTGATCCTGGCATCTGTGGTCTATGTAA

Vmn1r175: AGTACAAACATGTGCTCCACCTGCTTTCTGAGCACTTATCAGCTTGTCAC

**NICD:**

Olfr1006: AGGGAACATGTTGCTGGTTGTTTTAATCCGAATTGATTCTAGACTGCATA

Tas2r105: GACCTCGGAGATGTACTGGGAGAAAAGGCAATTCACTATTAACTACGTTT

Vmn1r139: AAGCATTGGCAAGTCACAGGCAAAGAGTGACACAGAGACGTTCCTCAATT

**VEGFA:**

Olfr1002: AGGCCTTATAAGCACTGTGGTCCATACTACTTCTGCATTTATTCTTCCAT

Tas2r104: TAACGTGGCTAGCTTCCTTTCCGCTAGCTGTGAAGGTCATTAAAGATGTT

Vmn1r12: ACTACATTGTCAGGAGCTTGATTTTAACTGTGACAACTTCCAGGGATATG

**shPTEN:**

Olfr1013: GTACACATTGACTTTGATGGGAAATAGCTCCCTCATTATGTTAATCTGCA

Tas2r110: ACTAGTGAATATCATGGACTGGACCAAGAGAAGAAGCATTTCATCAGCGG

Vmn1r170: TGATTCTCCTGAACAGACACCACCACAGACTGCAGCATATTCAATCCACA

Olfr1015: TGTCTATGTGAAAATCCTTTCCAGTATGGTGGGCTTCACTGTCCTCTCAA

Tas2r114: TGTAATTTGTCTGTTAATCCCAGAAAGCAACTTGTTATTCATGTTTGGTT

Vmn1r172: GGAAGTAAATGCCCAGAGAGTCTTCAAAGGAAGACAGTCATAGCTGTTTT

**shREN:**

Olfr1008: CCAGGCTCTGCTATTCACCAGTAAAATTTTCACATTAACTTTCTGTGGCT

Tas2r106: AAGGCACTGAAGCAATTAAAATGCCATAAGAAAGACAAGGACGTCAGAGT

Vmn1r157: CAGATCCTCTTGCTTTGCCATTTTGAGGTTGGGACCGTGGCCAATGTCTT

**shTRP53:**

Olfr1009: CCAGAGACTCTGCATACAGCTGGTGATCGGACCCTATGCTGTTGGCTTTT

Tas2r107: GCTCTCTAAGATCGGTTTCATTCTCATTGGCTTGGCGATTTCCAGAATTG

Vmn1r167: GTTTCAGTATAGGCATGCGCATCTTATCATTTGCCCATGATGGAGTGTTC

Olfr1014: TTGCTGTGTATGCATTAACTGTGTTAGGAAACAGCACCCTCATTGTGTTG

Tas2r113: GATCAATCATTGTAACTTTTGGCTTACTGCAAACTTGAGCATCCTTTATT

Vmn1r171: AACAGCACTGCCCTCATGATCACTATTCCGTTGACCAATGAAGTTGTCTC

Olfr107:  TTACTGCTTTCTTGCTCAGACACTCACCTCAGTGAGGGCCTGATGATGGC

**shKMT2C:**

Olfr1012: ATCTACTCTCGGCCAAGTTCCAGTTATTCCTTGGAAAGGATAAAATGGT

Tas2r109: TTCTAGAATTTTCCTGCTCTGGTTCATGCTAGTAGGTTTTCCAATTAGCT

Vmn1r169: GGTACCTGGGGTAGGGTGATGCTCCATGGAAGAGCCCCCAAATTTGTGAG

#### Histopathology

After fixation, representative specimens of the liver were routinely dehydrated, embedded in paraffin, and cut into 4 µm-thick sections. Tissue sections were stained with hematoxylin and eosin (H&E) according to standard protocols. Slides were scanned using a SCN400 slide scanner (Leica Biosystems) at 20X magnification.

Nodule annotation was initially performed by experienced pathologists (D.F.T, H.W.) based on H&E-stained sections using quPath software and 10X loupe browser software. Nodule annotation was further refined based on specific transcript expression (Extended Data Fig. 5) using 10X loupe browser software (H.W., M.B.).

#### Immunohistochemistry

After heat-induced antigen retrieval at pH6 or pH9, FFPE tissue sections were incubated overnight with the primary antibody and blocked with hydrogen peroxide if necessary. Depending on the primary antibody, an anti-mouse or anti-rabbit secondary antibody conjugated to HRP and AP, respectively, was applied (PolyviewPlus, ENZO Life Sciences GmbH, Lörrach, Germany). The signal was visualized using either 3,3'-Diaminobenzidine (Dako Liquid DAB+ Substrate, Agilent Technologies, Inc., Santa Clara, USA) or alkaline phosphatase (Permanent AP Red, Zytomed Systems, Berlin, Germany) as a chromogen. Details are given in Methods Table 1.

| Antibody | Host | Company | Order number | Antigen retrieval | Dilution | Detection reagent | Chromogen | Blocking |
|---|---|---|---|---|---|---|---|---|
| CK19 | Rabbit | Abcam | ab 133496 | Dako Target Retrieval Solution 10X Concentrat | 1:100 | PolyviewPlus AP anti rabbit | Permanent AP Red | / |

| | | | | | | | | |
|---|---|---|---|---|---|---|---|---|
| | | | | e, pH9, REF S2367 | | | | |
| HNF4alpha | Rabbit | Abcam | ab181604 | Dako Target Retrieval Solution 10X Concentrate, Citrate pH6, REF S2369 | 1:400 | PolyviewPlus AP anti rabbit | Permanent AP Red | / |
| GS | Mouse | BioScience | BD610517 | Dako Target Retrieval Solution 10X Concentrate, Citrate pH6, REF S2369 | 1:1000 | PolyviewPlus HRP anti mouse | DAB | $H_2O_2$ |
| tRFP | Rabbit | Evrogen | AB 233 | Dako Target Retrieval Solution 10X Concentrate, Citrate pH6, REF S2369 | 1:500 | PolyviewPlus AP anti rabbit | Permanent AP Red | / |
| GFP | Rabbit | Cell Signaling | 2956 | Dako Target Retrieval Solution 10X Concentrate, pH6, REF S1699 | 1:100 | PolyviewPlus HRP anti rabbit | DAB | $H_2O_2$ |

Table 1. Accompanying information related to immunohistochemistry.

Slides were scanned using a SCN400 slide scanner (Leica Biosystems) at 20X magnification. The individual histochemical GFP, RFP, and GS staining was evaluated as high = very intense, uniform staining, moderate = moderate intensity or intense non-uniform staining, low = low intensity and non-uniform staining using quPath software. Mapping of barcode signals to respective IHC data was performed by manual assessment of marker staining on IHC images using quPath software (H.W., M.B.). Next, corresponding tumor nodules were selected, categorized and stratified using 10X loupe browser software (H.W., M.B.). Note that this approach can be error-prone due to shifts in z-plane based on serial sectioning for each individual IHC sample and samples used for 10X Visium.

**Genotyping**

### Barcode expression pre-preprocessing

Before analyzing the 10X Visium data, we applied a filtering criterion of UMI counts > 5000. For the CytAssist platform, we excluded the outermost layer of spots due to unexpectedly high UMI counts. In addition to manually identifying cancerous nodule regions, we also annotated "normal tissue" regions to acquire representation of areas without any cancerous cells. The selection of normal regions was based on a minimum distance of 250-700 μm from the nearest tumor, depending on the tissue section, to minimize contamination from adjacent tumor regions. Tumor nodules were defined as described in the histopathology methods section.

### Bayesian modeling of perturbation probabilities from barcode counts

In our model, the observed expression count matrix $D_{s,b}$ (spots $s$ by barcode genes $b$) is assumed to follow a Negative Binomial distribution. This matrix has a mean $\lambda_{s,b}$ and overdispersion $\phi_b$. The overdispersion parameter $\phi$ is sampled from a Gamma distribution (shape=1000, rate=0.03), skewed towards higher values to encourage the likelihood to approximate a Poisson distribution in the absence of overdispersion evidence.

The mean expression for each spot, $\lambda_{s,b}$, is calculated as:

$$\lambda_{s,g} = \mu_s \sum_r A_{s,r} \sum_g G_{r,g} B_{g,b} k_b + \xi_b$$

where $\mu_s$ represents the sensitivity of each spot, $A_{s,r}$ maps spots to clonal nodules $r$, $G_{r,g}$ estimates the expected number of integrated copies of plasmid $g$, $B_{g,b}$ links plasmids to their corresponding barcodes, and $k_b$ is the barcode expression rate. $\xi_g$ is a barcode-specific additive noise term.

The per-nodule plasmid integration number, $G_{r,g}$, is modeled as an expected count of integration events, described by $F_{r,g,o}$. Here $F_{r,g,0}$ captures the probability of no integration, and higher indices reflect the integration of increasing numbers of copies. This is modeled using a Dirichlet distribution with a uniform concentration parameter and an order $o$ = 6. This assumes the maximum of six copies of the same plasmid per clone, balancing the need to capture dosage-dependent variation with the practicality of limiting the number of parameters to be learnt. For "normal tissue" regions the probability of perturbation presence was fixed to $10^{-3}$ to indicate near absence, but not zero, in order to prevent numerical instabilities.

The barcode expression rate, $k_g$ as well as additive noise $\xi_g$ , are sampled from a weakly regularized Exponential distribution with a rate of 1. Spot sensitivity $\mu_s$ is modeled by a Gamma distribution (shape=3, rate=0.3) that is weakly centered around 1. This parameter accounts for both the sensitivity variability across 10x Visium spots and the dilution effects on the barcode signal due to varying tumor purity.

**Perturbation probability model inference**

We infer our Bayesian model using a variational posterior approximation. Specifically, we employ a Log-Normal guide distribution to approximate the parameters that have Exponential and Gamma distributed priors. Additionally, we use a Dirichlet approximation for the posterior of $F_{r,g,o}$. The model and its inference framework are implemented in Pyro (v1.8.6) [14]. The variational approximation is conducted via the Stochastic Variational Inference (SVI) method [15], employing the Adam optimizer set at a learning rate of 0.01 and using 3 samples for KL divergence estimation. We perform inference over 10,000 gradient steps, monitoring the Evidence Lower Bound (ELBO) to assess convergence.

**Occurrence of individual perturbation combinations**

Considering the probabilistic nature of our estimates for $F_{r,g,o}$, we utilized samples from the estimated posterior to analyze tumor populations. Due to uncertain integration copy number estimates, we focused on presence/absence categories. These probabilities were computed as $1 - F_{r,g,0}$, and representative genotypes were sampled with Bernoulli distribution for each region and perturbation. We aggregated the data across 322 nodules into 256 possible genotype states, which allows us to compute medians and confidence intervals for marginal integration numbers (indicating the count of different plasmids integrated) and individual perturbation for frequencies, as well as frequencies of individual genotypes.

While it may be tempting to interpret high frequencies of genotype occurrence as advantageous for tumor proliferation, such raw frequencies could be confounded by technical factors, such as initial plasmid concentration and integration rate. To address this, we constructed a null hypothesis (H0) over the 256 individual genotype numbers.

This hypothesis holds the marginal expected number of integrations and perturbation frequencies constant across the population, attributing variations solely to technical effects, and assumes that the genotypes are independently distributed.

In practice, we adjust the observed perturbation frequencies to account for technical biases by normalizing these frequencies - dividing the average observed perturbation frequency for each perturbation by the sum of all plasmid frequencies and multiplying by the expected number of integrations. We then simulate the distribution of genotypes by drawing Bernoulli samples using these rescaled probabilities for each perturbation. This process is repeated for the number of nodules (322), and the results are aggregated back into the 256 genotype states to create a sampling strategy that reflects the desired properties. By comparing deviations between 5000 samples drawn from both the inferred posterior (Observed) and the simulated H0 (Expected), we can identify genotypes with significant tumorigenic effects (Observed > Expected) or disadvantageous effects (Observed < Expected).

**Co-occurrence odds ratios and model of second order interaction effect**

With posterior estimates of the genotypes within the tumor population, we can test for interaction effects between individual perturbations. We categorize the frequencies of perturbations A and B into four groups: $p_{00}$ (A-/B-), $p_{01}$(A-/B+), $p_{10}$ (A+/B-), and $p_{11}$ (A+/B+). The system can be described using a softmax linear model expressed as:

$$p_{i,j} = \frac{\exp(\theta_{00}+i\theta_{10}+j\theta_{01}+ij\theta_{11})}{\sum_{ij}\exp(\theta_{00}+i\theta_{10}+j\theta_{01}+ij\theta_{11})}$$

Here, $\theta_{10}$ and $\theta_{01}$ represent the effects of individual perturbations, and $\theta_{11}$ is the interaction effect. By setting $\theta_{00}$ to zero to eliminate softmax non-identifiability and using $Z$ as the normalization constant $\sum_{i,j} e^{\theta_{i,j}}$, we derive the following relationships:

$$p_{10}/p_{00} = [e^{\theta_{10}}/Z]/[1/Z] = e^{\theta_{10}}$$

Similarly,

$$p_{01}/p_{00} = e^{\theta_{01}}$$

and

$$p_{11}/p_{00} = e^{\theta_{10}+\theta_{01}+\theta_{11}}$$

Thus, computing the pairwise odds ratios (OR) effectively determines the interaction effect $\theta_{11}$:

$$OR = \frac{p_{11}p_{00}}{p_{10}p_{01}} = \frac{p_{11}/p_{00}}{[p_{10}/p_{00}][p_{01}/p_{00}]} = \frac{e^{\theta_{10}+\theta_{01}+\theta_{11}}}{e^{\theta_{10}}e^{\theta_{01}}} = e^{\theta_{11}}, \blacksquare$$

For each gene pair, we estimated the ORs and assessed their significance by drawing 2000 samples from the posterior probability of perturbation presence for each nodule. By fitting a softmax linear model directly to the data and setting the interaction effect $\theta_{11}$ to zero (OR = 1), we simulated the expected probabilities for the pairwise groups under the assumption of no interaction.

**Genotype to phenotype GLM**

To explore the relationships between inferred perturbation probabilities and phenotypic features — specifically, IHC staining status and gene expression — we employed a generalized linear model (GLM). Here, the inferred perturbation probabilities serve as the explanatory variables $X$.

For the IHC staining analysis conducted at the nodule level, we used binary annotations (positive/negative) and modeled the outcomes with a Bernoulli distribution. The staining status for each nodule, $Y_{r,m,k}$ ($r$ - region, $m$ - gene, $k$ - sample), is modeled as:

$$Y_{r,m,k} \sim Bernoulli(\sigma(\sum_g X_{r,g} w_{g,m}) + z_k)$$

where $\sigma(x) = \frac{1}{1+e^{-x}}$ is the sigmoid link function. The weight matrix $w_{g,m}$, akin to L1 regularization, is sampled from a Laplace distribution centered at zero with a scale parameter $b$. We set the scale to 1 for the intercept and 0.1 for the perturbation weights to impose stronger regularization on the perturbations. The batch effect $z_k$ for each sample $k$ is also sampled from a Laplace distribution (0, 1). The explanatory variable $X_{r,g}$ is directly sampled from $1 - F_{r,g,0}$, estimated by the perturbation probability model (non-learnable in the GLM).

Gene expression is modeled similarly, with few key differences. As gene expression is recorded as a nonzero integer at the spot level $s$, we use a Poisson distribution:

$$Y_{s,m,k} \sim Poisson(\mu_s \exp(\sum_g X_{s,g} w_{g,m} + z_k))$$

In addition to the parameters used in the IHC model, spot sensitivity $\mu_s$ is factored in, sampled from the posterior of the perturbation probability model. $X_{s,g}$ is calculated as $\sum_r A_{s,r}(1 - F_{r,g,0})$. The weight matrix $w_{g,m}$ for perturbation-related weights is sampled from a Laplace distribution centered at zero with a strongly regularizing scale $b = 10^{-3}$.

#### GLM Inference

We infer our Bayesian model using a mean field variational posterior approximation. The model and its inference framework are implemented in Pyro (v1.8.6)[14]. The variational approximation is conducted via the Stochastic Variational Inference (SVI) method[15] , employing the Adam optimizer set at a learning rate of 0.01 and using 3 samples for KL divergence estimation. At each gradient descent step, the parameters $X$ and $\mu$ are sampled from their respective posterior distributions as estimated by the perturbation probability model. This approach integrates the uncertainties associated with their estimation directly into the GLM framework. We perform inference over 2,000 gradient steps, monitoring the Evidence Lower Bound (ELBO) to assess convergence.

#### Reading Visium space ranger output into data objects

We utilized the anndata package (v0.11) in Python and the SingleCellExperiment package (v1.24.0) in R to generate and manage 10X Visium data objects. To facilitate communication between R and Python, we employed ZellKonverter (v1.12.1). For data processing, we employed Scanpy (v1.9.8)[16], squidpy (v1.4.1)[17], and scater (v1.30.1)[18] in Python and R. We refined our analysis by subsetting all objects to include only features shared across all 11 slides, resulting in a total of 19,464 genes.

#### Publicly available databases of cell type markers

We used scLiverDB[19] , PanglaoDB[20] , and MSigDB[21] to collect an initial set of marker genes for prevalent cell types and gene sets in normal and tumor liver tissues of mouse and human. This yielded a list of a total 2323 genes.

#### Data normalization and preprocessing

After applying filtering criteria to exclude genes with raw counts less than $10$ or greater than $10^6$ for any single slide, as well as barcode genes, the count matrices were normalized for each spot to ensure a total count of $10^4$. Subsequently, the normalized values were log-transformed ($log(x+1)$). This preprocessing was executed in Python using Scanpy (v1.9.8). Utilizing Scanpy (v1.9.8) with the Seurat flavor, we identified $15,000$ highly variable genes for each of the $11$ Visium and Visium Cytassist slides. The intersection of these sets resulted in $9,205$ genes. Subsequently, in the final refinement step, we narrowed down the gene set to $80$ core markers, resulting in $7,361$ genes. This final gene set was employed for all subsequent phenotype analyses and visualizations.

**Gene co-expression networks**

Using the spot-level normalized expression values after filtering out uninformative genes, we conducted Gaussian graphical modeling (GGM)[22] to infer a sparse gene co-expression network. We utilized the R package glasso (v1.11) for this purpose. The regularization parameter was optimized through a grid-search approach and set to $0.3$. Following the construction of the initial GGM, we refined the network by filtering edges to retain only those with a Pearson's pairwise correlation coefficient of at least $0.25$. The isolated genes are, in turn, dropped from the graph. We used the R package igraph (v2.0.1.1) to visualize the graphs.

**Nodule-level expression aggregation**

We computed two types of aggregates for normalized expression values within each nodule: mean-based and quantile-based. For the mean-based aggregates, we calculated the average normalized expression of each gene across all spots within each nodule. For the quantile-based aggregates, we determined $q_{95}$ of expression values across all spots per nodule. We used these aggregates in the subsequent nodule-level analyses.

**Binarization of nodule phenotypes**

After quantile-normalization we computed the average of the scaled expression values for the core markers per phenotype, we then thresholded the values by 0.5, where all nodules with the aggregate value above 0.5 are considered to have the corresponding phenotype signature and otherwise not. The mean-based aggregates for each gene are quantile-normalized further by $\frac{x - q_{25}}{q_{99} - q_{25}}$ where $q_{25}$ and $q_{99}$ represent the $25$th and

$99$th quantile values of the aggregate gene expression for the corresponding gene across all nodules and all slides. We then binarized the values above $0.5$ as $1$ (on) and otherwise $0$ (off).

#### Binarization of nodule TME signatures

We binarized the TME signatures following the same procedure as for nodule phenotypes, but using the initial quantile-based aggregates instead.

#### Phenotype and TME heatmaps

We generated heatmaps of scaled gene expression using the processed expression values at the nodule level, employing ComplexHeatmap (v2.16.0)[23]. Clustering of both rows and columns was performed based on Spearman's correlations. Additionally, hierarchical clustering was applied to subdivide genes into clusters. The color bar associated with genes indicates their corresponding phenotypes, with emphasis on the core markers. An attached annotation heatmap illustrates scaled ($p^{10}$) estimated plasmid probabilities per nodule.

#### Spatial integration and nodule unification

To integrate all slides into a unified embedding space and classify spots based on their phenotypic signatures in an unbiased manner, we employed an ensemble spatially-aware classifier implemented in SageNet (v1.1.0)[24]. Data from all slides were trained and fed into this classifier. Subsequently, we clustered the spots within the embedded space using Scanpy's wrapper of Leiden clustering (with a resolution of 1)[25]. We then performed voting classification to classify nodules to the most dominant class across the spots belonging to the corresponding nodules. We call these classes the "unified nodule annotations". Finally, we extracted spatially informative genes from the SageNet model.

#### Inter-nodule differential gene expression

To delve deeper into inter-nodule transcriptional differences, we conducted differential gene expression analysis using the FindMarkers method from the R package scran (v1.28.2). This method allowed us to perform a light-weight differential gene expression analysis on the unified nodule annotations.

#### Celltype-informed Factor Analysis

We concatenated all raw anndata objects and subsetted it to the set of "core markers" and associated genes (as listed in Fig. 3b and Fig. 3d) as well as $500$ highly variable genes across slides. We then used cell2module (github.com/vitkl/cell2module) to perform Non-negative matrix factorization. Cell2module model treats raw RNA count data $D$ as negative-binomial distributed, given transcription rate $\mu_{c,g}$ and a range of variables accounting for technical effects:

$$D_{c,g} \sim \texttt{NB}(\mu = \mu_{c,g}, \alpha_{a,g})$$

$$\mu_{c,g} = ((\sum_f w_{c,f} g_{f,g}) + s_{e,g}) * y_c$$

Where $\mu_{c,g}$ denotes expected RNA count $g$ in each cell $c$. $\alpha_{a,g}$ denotes per gene $g$ stochastic/unexplained overdispersion for each covariate $a$; $w_{c,f}$ denotes cell loadings of each factor $f$ for each cell $c$; $g_{f,g}$ denotes gene loadings of each factor $f$ for each gene $g$; $s_{e,g}$ denotes additive background for each gene $g$ and for each experiment $e$, to account for contaminating RNA; $y_c$ denotes normalization for each spot $c$, to account for RNA detection sensitivity, sequencing depth. We recovered $40$ factors representing groups of coexpressing cell type signatures using the default cell2module parameters. After training, we inferred the posterior of the gene loadings per factor. Subsequently, we extracted genes with the top $5$ posterior median values and compared them to predefined marker gene lists per cell type. Finally, we mapped each factor to the cell type with the most number of overlapping genes.

### REFERENCES_METHODS

23. Gu, Z., Eils, R. & Schlesner, M. Complex heatmaps reveal patterns and correlations in

multidimensional genomic data. *Bioinforma. Oxf. Engl.* **32**, 2847–2849 (2016).

24. Elyas Heidari *et al.* Supervised spatial inference of dissociated single-cell data with SageNet. *bioRxiv* 2022.04.14.488419 (2022) doi:10.1101/2022.04.14.488419.

25. Traag, V. A., Waltman, L. & van Eck, N. J. From Louvain to Leiden: guaranteeing well-connected communities. *Sci. Rep.* **9**, 5233 (2019).
